# HLA topography enforces shared and convergent immunodominant B cell and antibody alloresponses in transplant recipients

**DOI:** 10.1101/2023.03.31.534734

**Authors:** John T. Killian, R. Glenn King, Aaron C.K. Lucander, James L. Kizziah, Christopher F. Fucile, Ruben Diaz-Avalos, Shihong Qiu, Aaron Silva-Sanchez, Betty J. Mousseau, Kevin J. Macon, Amanda R. Callahan, Guang Yang, M. Emon Hossain, Jobaida Akther, Daryl B. Good, Susan Kelso, Julie A. Houp, Frida Rosenblum, Paige M. Porrett, Song C. Ong, Vineeta Kumar, Erica Ollmann Saphire, John F. Kearney, Troy D. Randall, Alexander F. Rosenberg, Todd J. Green, Frances E. Lund

**Affiliations:** Department of Surgery, at The University of Alabama at Birmingham Heersink School of Medicine, Birmingham, AL 35294, USA; Department of Microbiology, at The University of Alabama at Birmingham Heersink School of Medicine, Birmingham, AL 35294, USA; Department of Informatics and Data Science, at The University of Alabama at Birmingham Heersink School of Medicine, Birmingham, AL 35294, USA; Center for Infectious Disease and Vaccine Discovery, La Jolla Institute for Immunology, La Jolla, CA 92037, USA; Department of Medicine: Divisions of Clinical Immunology and Rheumatology, at The University of Alabama at Birmingham Heersink School of Medicine, Birmingham, AL 35294, USA; Nicoya, Kitchener-Waterloo, Ontario, Canada; Department of Pathology, at The University of Alabama at Birmingham Heersink School of Medicine, Birmingham, AL 35294, USA; Nephrology, at The University of Alabama at Birmingham Heersink School of Medicine, Birmingham, AL 35294, USA

**Keywords:** B lymphocytes, Antibodies, Alloimmunity, Transplantation, Epitopes, Immunodominance, Topography

## Abstract

Donor-specific antibody (DSA) responses against human leukocyte antigen (HLA) proteins mismatched between kidney transplant donors and recipients cause allograft loss. The rules governing the immunogenicity of non-self donor HLA are poorly understood. Using single-cell, molecular, structural, and proteomic techniques, we profiled the HLA-specific B cell response in the kidney and blood of a transplant recipient with antibody-mediated rejection (AMR). We observed an immunodominant B cell antibody response focused on topographically exposed, solvent-accessible mismatched HLA residues along the peptide-binding groove – a subregion comprising only 20% of the HLA molecule. We further demonstrated that, even within a diverse cohort of transplant recipients, the B cell alloresponse consistently converges on this same immunodominant subregion on the crown of the HLA molecule. Based on these findings, we propose that B cell immunodominance in transplant rejection relies on antigenic topography, and we suggest that this link could be exploited for organ matching and therapeutics.

## INTRODUCTION

Antibodies (Abs) against mismatched human leukocyte antigens (HLA) are a leading cause of injury and graft loss in kidney transplantation.^1–3^ Up to 25% of kidney transplant recipients will develop a new donor-specific anti-HLA antibody (HLA-Ab) following transplant,^4^ and these individuals suffer a 40% absolute decline in 10-year graft survival.^5^ Ultimately, graft loss and the re-initiation of dialysis is associated with significant increases in mortality for these transplant recipients.^6^ In addition to antibody-secreting cells (ASCs), which produce injurious donor-specific Abs (DSA), the presence of donor HLA-specific (alloreactive) circulating memory B cells (Bmem) also correlates with poor outcomes.^7–9^ Thus, both arms of the humoral immune response work in concert to drive rejection.^10^ Unfortunately, current therapies aimed at managing antibody-mediated rejection (AMR) and depleting ASCs and Bmem are largely ineffective in transplant rejection, indicating the unmet need to both prevent and treat these alloresponses.^11^

To produce high-affinity Abs against a target antigen, the B cell response relies on the competitive environment of the germinal center (GC) within secondary lymphoid tissues where the processes of somatic hypermutation (SHM) and clonal selection result in the development of expanded, affinity-matured lineages of long-lived ASCs and Bmem.^12^ Studies of infection and vaccination show that the quality of lymphoid tissue GC-driven responses is regulated by the assistance of other immune cells,^13^ like T follicular helper cells (T_FH_ cells), as well as features of the antigen.^13^ For example, B cell responses are influenced by the mode of antigen exposure, the persistence of antigen and the physical characteristics of the antigen.^14^ Individual B cells respond to protein antigens by recognizing exposed solvent accessible epitopes.^15^ In many cases, B cells can target multiple distinct accessible epitopes within a single antigen, resulting in a diverse polyclonal repertoire of Bmems and ASCs – all of which are specific for the same antigen.^16^ For other antigens, the B cell response may be more tightly focused on a limited number of immunogenic epitopes at the expense of other epitopic sites. This focused “immunodominant” B cell response, chiefly described in anti-viral immunity, is regulated by the frequency of epitope-specific B cells, prior antigen history (memory), affinity, antigen topography, the duration of antigen delivery, and T cell help.^17,18^ In addition, single polymorphic residues with the antigen may contribute to immune focusing and immunodominance across B cell and ASC compartments.^19,20^

Although HLA residues that are mismatched between donor and recipient are necessary to generate alloimmunity^21^ and clearly lead to the production of HLA-Abs, AMR, and graft loss,^22^ the relative immunogenicity of individual polymorphic residues and the rules governing immune responses to HLA antigens remain largely unknown.^23–25^ This lack of knowledge is compounded by the fact that alloreactive B cell responses are difficult to sample^26^ as organ-infiltrating alloreactive B cells^27^ and tertiary lymphoid structures can be found in transplanted organs.^28,29^ Moreover, alloreactive B cell responses focus on an antigen, HLA, that is persistent, abundant, and quite similar at a structural and molecular level to self-HLA. One widely used model for interpreting studies of HLA-Ab responses and understanding what mismatches constitute “self” vs. “non-self,” posits that small (3.5 Å-radius) patches of mismatched polymorphic residues within the larger HLA epitope, so called eplets, are the critical determinants of the HLA-specific B cell response.^30^ However, there are no structural data to support this model. In fact, the only reported HLA-Ab/HLA complex structure^31^ features an Ab that was isolated using phage display, rather than an Ab that was generated during an ongoing alloimmune response in a transplanted individual.

Given this knowledge gap, the United States Kidney Allocation system currently prioritizes only perfect whole-antigen matches between donor and recipient^32^ – an approach that converts the matching problem into an “all-or-nothing” proposition. Given the evidence showing that increasing numbers of amino acid (AA) mismatches between donor and self-HLA correspond with poor transplant outcomes,^22,33–35^ there is considerable interest in matching donors and recipients at the residue level.^23,36–39^ To take this approach, it is critical to first understand the rules governing allorecognition by B cells. In this manuscript, we used single-cell, molecular, structural, and proteomic techniques to assess the alloimmune response in a kidney transplant recipient (subject N006) with clinical features of anti-HLA DSA, AMR, and graft loss. Kidney and blood B cells isolated from recipient N006 revealed that alloreactive HLA-A*01:01-specific B cells underwent clonal selection and affinity maturation. This high-affinity alloreactive B cell response not only spanned B cells and ASCs in the rejected allograft and the peripheral blood but also included lineages of Bmems and ASCs that were clonally linked to circulating A*01:01-specific DSA in the individual. Profiling the B cell response against a mismatched donor HLA protein at the single cell level revealed multiple alloreactive A*01:01-specific B cell clonal lineages but also clearly showed epitopic focusing and immunodominance of the allo-HLA B cell response. We determined the structure of three Ab/A*01:01 HLA complexes and revealed the structural basis of these immunodominant HLA epitopes. Using competitive Ab binning and single-residue HLA mutants, we demonstrated that the B cell and serologic alloresponses of this individual converged on the topographically exposed “crown” of HLA-A*01:01. Finally, we examined a larger cohort of transplant patients who generated a *de novo* A*01:01 DSA after receiving a mismatched A*01:01 organ. Although these individuals all expressed unique combinations of self-HLA-A molecules, their A*01:01 DSA response converged on the same immunodominant, crown-localized HLA-A*01:01 epitopes that we identified in our first transplant subject. Together, these data show B cell alloimmune responses are highly focused and directed against a limited number of non-self residues localized within a discrete small topographic region in A*01:01 – a “public” region that is conserved across individuals expressing diverse constellations of self-HLA-A proteins. Thus, while mismatched residues in alloAb responses to HLA-A are clearly important in driving transplant rejection, the data suggest that the most immunogenic mismatched residues may be restricted to the small number of mismatched residues that are localized within this specific topographically defined region of the HLA antigen.

## RESULTS

### Transplant recipient kidney and blood B lineage cells display features of antigen-driven selection

ASCs that produce Abs directed against HLA molecules that are mismatched between the allograft donor and recipient (donor-specific Abs or DSA) are important drivers of chronic kidney allograft rejection,^1–3^ yet we know remarkably little about the structural epitopes in the mismatched HLA molecules that are specifically targeted by the humoral immune response.^25,31^ Likewise, the relationship between the recirculating or allograft tissue-residing alloreactive B cells and the donor HLA-specific ASCs and Abs is not well-characterized.^27^ To evaluate the contribution of alloreactive B cells to allograft rejection, we examined the tissue and circulating donor HLA-specific B lineage response in a kidney allograft recipient (subject N006) who presented with histopathologic allograft injury consistent with AMR and expressed high levels of circulating DSA directed against the mismatched HLA allele (HLA-A*01:01) expressed on the donor kidney (Table S1).

Following surgical nephrectomy to remove the rejected kidney from recipient N006, we performed immunohistologic and flow cytometric analyses on tissue sections and cells isolated from the allograft. Although B cells have been reported to co-localize with T_FH_ and follicular dendritic cell networks in GC-like structures^28^ within the graft, we did not identify tertiary lymphoid structures in this allograft (Fig. 1A-C). However, we observed clusters containing T and B cells (Figure 1A), proliferating B cells (Figure 1B), and ASCs (Figure 1C). Furthermore, we identified multiple B cell subsets, including IgD^neg^CD27^+^ Bmem, IgD^neg^CD27^neg^ (double negative, DN) B cells, and CD27^hi^CD38^hi^ ASCs (Figure 1D), in the graft and observed that many of these intragraft B cells expressed the Bmem activation marker CD71 (Figure 1E). Thus, the allograft contained activated, proliferating antigen-experienced B cells and ASCs.

**Figure 1.**
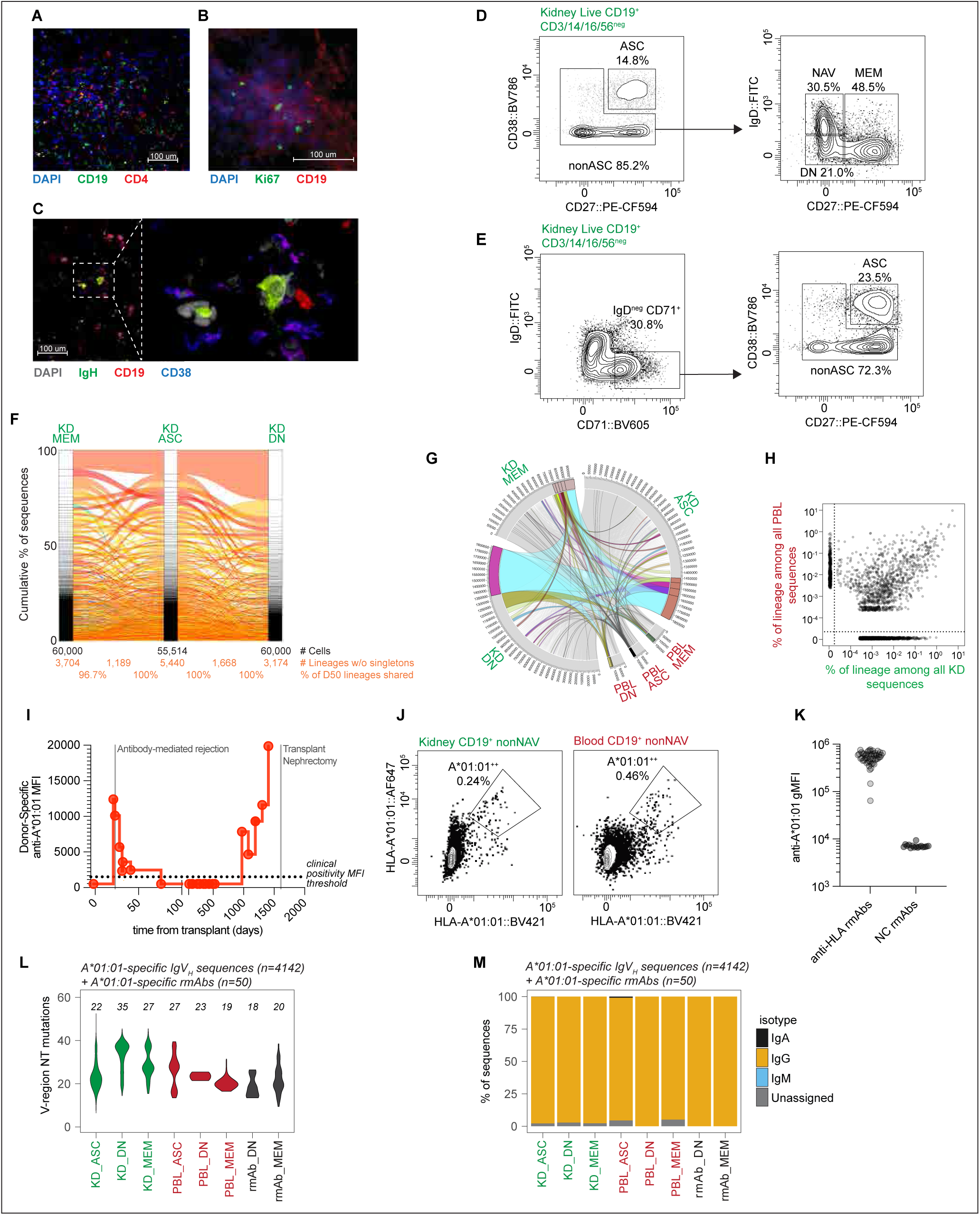
Somatically mutated, class-switched alloHLA-specific memory and effector B cells are found in circulation and in the allograft of a kidney transplant recipient. **(A-C)** Immunofluorescent histology of a rejected kidney (KD) allograft from recipient N006 showing CD4^+^ T (red) and CD19^+^ (green) B cells **(A)**, proliferating Ki67^+^ (green) CD19^+^ (red) B cells **(B)**, and CD19^+^ (red), CD38^+^ (blue), intracellular IgH^+^ (green) ASCs **(C)**. DAPI nuclear staining shown as blue in **(A)** and **(B)** and gray in **(C)**. **(D-E)** Flow cytometry analysis of recipient N006 kidney allograft to identify dump negative (CD3/CD14/CD56^neg^) B lineage (CD19^+^) subsets including ASCs (CD27^+^CD38^+^), naïve (NAV, CD27^neg^IgD^+^), Bmem (MEM, CD27^+^IgD^neg^) and CD27^neg^IgD^neg^ DN B cells (**D**) and CD71 (**E**) to identify activated cells. **(F)** Alluvial plots showing lineage relatedness between ASCs, DN B cells, and Bmem cells isolated from recipient N006 KD allograft. Shared lineages (ribbons) ranked by size in each B cell subset with lineages shared between all 3 subsets (orange) or between any two subsets (green). Numbers of sorted cells and non-singleton lineages for each subset provided. %D50 defined as: (# lineages in top 50% of subset A / # of lineages in the top 50% of subset B) × 100. See *Methods* for details. **(F)** Circos plot showing connections between lineages of B cell subsets isolated from recipient N006 KD allograft and peripheral blood (PBL). Size-ranked lineages representing the top 20% of size-ranked sequences within a given subset are colorized. See *Methods* for details. **(G)** Scatterplot comparing size of B cell lineages in recipient N006 KD (x-axis) and PBL (y-axis). Lineage size shown as % sequences of that lineage within KD or PBL. Lineages not detected in a tissue reported as 0%. **(I)** Trajectory of anti-HLA-A*01:01 DSA response in recipient N006 based upon clinical HLA-Ab testing. Clinical index diagnosis of AMR and timing of transplant nephrectomy indicated. **(I)** Fluorochrome-labeled recombinant HLA-A*01:01 tetramer binding to CD19^+^IgD^neg^ B cells isolated from recipient N006 kidney and blood. Dual tetramer-stained cells (gated population) were index-sorted for single cell rmAb production. **(J)** Binding to HLA-A*01:01-coated microbeads by 50 rmAbs from single-cell sorted HLA-A*01:01-specific B cells (see **J**) or human rmAbs specific for influenza HA protein (negative control, (NC)). Data reported as geometric mean fluorescent intensity (gMFI). **(L-M)** Mutation frequencies (**L**) and isotype distribution (**M**) in KD and PBL B cell derived IgV_H_ sequences that were assigned to A*01:01-specific B cell lineages. Violin plots (**L**) showing IgV_H_-region nucleotide (NT) mutation frequency distributions with median number of mutations in each bulk subset and rmAb subset shown. See Supplemental Figures and Tables for: schematic of experimental design (Figure S1); clinical description of transplant recipient N006 (Table S1); analyses of bulk B cell subsets (Figure S1); analyses of HLA-A*01:01-specific B cells and A*01:01-specific rmAbs (Figure S2); sequence data for A*01:01-specific IgV_H_ lineages and rmAb (Table S2); and cytometric bead binding data for A*01:01-specific rmAbs (Table S3).

To address whether the B cells and ASCs found within the allograft were distinct from the systemic circulating pool of ASCs and B cells, we sort-purified bulk populations of ASCs and antigen-experienced Bmem and DN subsets from the blood and the kidney (Figure S1A,B) and sequenced the B cell receptor (BCR) Ig heavy (IgH) chain variable domain (IgV_H_) of these cells. We assigned the IgV_H_ sequences to clonal lineages, which we defined as sharing *IGHV* and *IGHJ* gene segments, having identical IgH chain complementarity determining region 3 (HCDR3) lengths, and exhibiting >85% nucleotide identity across the HCDR3.^40^ Then, we analyzed clonal relationships between the B cells and ASCs found in circulation and those in the allograft. We observed extensive clonal expansion of B lineage cells within the kidney allograft, with rich connections between the ASC, Bmem, and DN B cell subsets (Figure 1F). Moreover, and in contrast to the low frequency of shared clonal lineages between blood and tissues isolated from a non-transplanted individual (Figure S1C), we identified numerous shared lineages between the B cells in the graft and blood of recipient N006 (Figure 1G, Figure S1C). We further observed that most lineages of B cells and ASCs present in the kidney were also found in the blood (Figure 1H). Next, we assessed the isotype and SHM profile of the BCRs expressed by the kidney and blood B cells. Most B cells in both sites were class-switched (Figure S1D), and the B cells and ASCs from both sites had undergone extensive SHM (Figure S1E). Thus, the B cells in the rejected kidney and blood of a transplant recipient undergoing AMR were isotype-switched, clonally expanded, shared, and mutated – all consistent with an ongoing robust adaptive immune response that spanned the allograft and blood.

### Class-switched, mutated alloHLA-specific Bmem and ASCs accumulate in the allograft and blood

Although clonally expanded B cells in kidney allografts^41^ and intragraft B cell infiltrates^42,43^ correlate with poor transplant outcomes, the specificity and origin of these B cells is still not well understood. Since standard clinical lab assays revealed that DSA for the allograft-derived HLA-A*01:01 was readily detected in the blood of transplant recipient N006 (Figure 1I), we hypothesized that some of the allograft-infiltrating, clonally expanded and affinity-matured B cells would be specific for the mismatched A*01:01 donor HLA allele. To test this hypothesis, we used fluorochrome-labeled tetramers of recombinant HLA-A*01:01 to sort-purify single A*01:01-specific B cells from the allograft and the peripheral blood (Figure 1J, Figure S2A,B) and then cloned, sequenced, and expressed the BCR IgH and Ig light (IgL) chain genes as recombinant human IgG1 monoclonal Abs (rmAbs). Using A*01:01-coated microbeads, we confirmed that 50 rmAbs, which were cloned from sorted A*01:01-specific Bmem (43/50) and DN B cells (7/50) isolated from the peripheral blood and kidney (Table S2), bound to the mismatched A*01:01 donor HLA protein (Figure 1K, Table S3). Next, we used the IgV_H_ sequences from the 50 A*01:01-specific rmAbs to evaluate whether the rmAbs derived from the A*01:01-specific B cells were clonally related to any of the expanded B cell lineages we identified in the bulk kidney and blood IgV_H_-Seq dataset (Table S2). If one or more A*01:01-specific rmAbs shared *IGHV* and *IGHJ* gene segments, identical HCDR3 lengths, and >85% nucleotide identity across the HCDR3 with a clonal lineage identified in the bulk IgV_H_-Seq dataset, we assigned the rmAbs to this clonal lineage and classified the entire lineage as A*01:01-specific. Using this approach, we identified 14 distinct A*01:01-specific lineages that linked the 50 A*01:01-specific rmAbs and the clonally related BCR IgV_H_ sequences (n=4142 sequences) derived from the bulk kidney and blood Bmem, DN B cells and ASCs (Table S2). We observed that the A*01:01-specific clonal lineages had undergone extensive SHM (Figure 1L) and that Bmem, ASCs and DN cells in the A*01:01-specific lineages were each heavily mutated (Figure S2C,D). While the total blood IgV_H_-Seq repertoire included many IgA sequences (Figure S1D), this was not observed in the A*01:01-specific lineages, which showed almost exclusive IgG utilization across all subsets in both blood and kidney (Figure 1M). Thus, multiple lineages of IgG-switched and highly somatically mutated B cells, specific for the mismatched donor A*01:01, were present in the blood and rejected kidney.

### Clonal lineages of allospecific affinity-matured B cells are shared across kidney and blood

As expanded clonal lineages of B cells reflect the immunogenetic record of SHM and antigen selection,^44,45^ we used the SHM profile of the A*01:01-specific B cell lineages to evaluate the evolution of the alloresponse to donor HLA. We identified many examples of lineages that contained multiple B lineage subsets (Figure 2A-C) that were found in both kidney and blood. As expected, these B cells and ASCs exhibited increasing numbers of mutations within the clonal lineage tree (Figure 2A,B). Moreover, 8 of the 14 A*01:01-specific lineages were each individually responsible for at least 2.5% of IgV_H_ sequences in the entire A*01:01-specific IgV_H_-Seq repertoire (Figure 2D) and together, these 8 clonal lineages collectively accounted for 93.0% of all A*01:01-specific sequences in the IgV_H_-seq database. At least two different types of B lineage populations were found in 10/14 lineages (Figure 2E), with the most represented subsets including kidney ASC (present in 11 lineages) and kidney Bmem (present in 8 lineages, Figure 2E). Next, we used localized surface plasmon resonance (L-SPR, Figure 2F) to measure the binding affinities (K_D_) of the 50 rmAbs specific for A*01:01 (Table S4). We determined that the median K_D_ of binding to A*01:01 by the 50 rmAbs derived from Bmem and DN B cells was 2.1 nM (Figure 2G, Table S4). To confirm that this high-affinity binding was due to the accumulation of mutations within the BCR and was not germline-encoded, we synthesized rmAbs encoding the inferred unmutated common ancestors (UCAs) of 9 different A*01:01-reactive lineages and assayed binding of the UCA rmAbs to A*01:01. As shown in Figure 2H, the K_D_s of two mutated rmAbs, which were members of either Lin4 (rmAb D01) or Lin12 (rmAb L02), were between 100- to 1000-fold lower (i.e. higher affinity) than the K_D_s of the rmAbs encoding the inferred UCA for Lin4 and Lin12. Similarly, we observed significant (p<0.0001) increases in A*01:01 affinity (lower K_D_ values) for all 36 rmAbs present in 9 different lineages when compared to the UCA rmAbs for each of these nine lineages (Figure 2I, Table S4). Interestingly, the UCAs from 5/9 lineages showed no detectable binding to A*01:01 by L-SPR (Figure 2J, Table S4). These data support the conclusion that the somatically mutated A*01:01 donor-specific B cells have undergone affinity maturation and have been selected for high-affinity clones.

**Figure 2.**
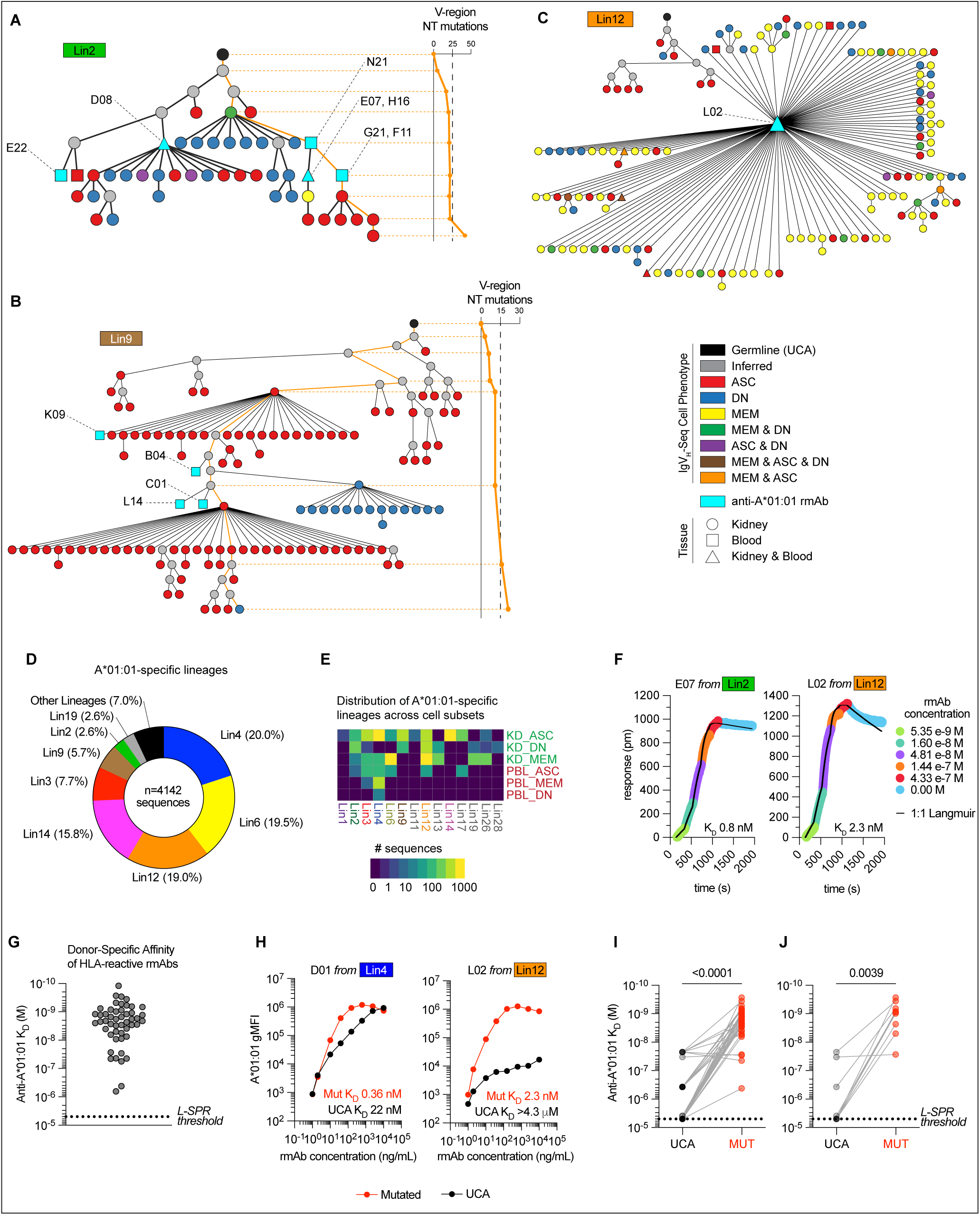
Identification of expanded clonal lineages of affinity-matured kidney and blood allo-HLA-A*01:01-specific B cells and ASCs. **(A-C)** Phylogenetic trees showing 3 representative IgV_H_ clonal lineages of A*01:01-specific B cells and ASCs. Tree edges trace IgV_H_ mutations from the inferred UCA (black node) through inferred mutation nodes (gray nodes) and observed nodes (colored symbol). Observed node color and shape denotes B cell subset(s) and tissue of origin assigned to each IgV_H_ node (see panel callout). Nodes containing IgV_H_ sequences from A*01:01-specific rmAbs indicated in cyan. Orange line indicates IgV_H_-region NT mutation accumulation. **(D)** Analysis of the A*01:01-specific IgV_H_-Seq repertoire (n=4142 sequences) from the bulk kidney and blood B cell subset IgV_H_ database described in Fig. 1F-H, 1L-M. Percentages of sequences assigned to the 14 different A*01:01-specific expanded lineages provided with callouts for individual clonal lineages representing ≥2.5% of the A*01:01-specific IgV_H_ sequence database. **(E)** Distribution of different B cell subsets within the 14 A*01:01-specific B cell clonal lineages. **(F)** Binding kinetics of A*01:01-specific rmAbs E07 and L02 to recombinant HLA-A*01:01. Binding reported as a function of increasing rmAb concentration, fit as 1:1 Langmuir curves by localized surface plasmon resonance (L-SPR). Affinity of binding reported as equilibrium constant (K_D_). **(G)** Affinities (K_D_) of 50 A*01:01-specific rmAbs binding to A*01:01. Dotted line indicates limit of detection (K_D_ 4.3 µM). **(H)** Binding to A*01:01 by representative Lin4 (rmAb D01) and Lin12 (rmAb L02) A*01:01-specific rmAbs and synthetic rmAbs produced using the sequences of the inferred Lin4 and Lin12 UCAs. Data shown as increasing concentrations of the rmAbs (x-axis) and gMFI of binding to A*01:01 cytometric beads (y-axis). L-SPR derived K_D_ values provided. **(I)** Paired analysis (n=36 pairs) showing L-SPR calculated K_D_ for A*01:01 by cloned A*01:01-specific rmAbs and the synthetic UCA rmAb that was predicted to be the non-mutated ancestor of each of the cloned somatically mutated rmAbs. **(J)** Paired analysis (n=9 pairs representing distinct A*01:01-specific B cell lineages) showing L-SPR calculated K_D_ for A*01:01 by cloned A*01:01-specific rmAbs (showing rmAb with the highest affinity in each lineage) and the synthetic UCA rmAbs for each lineage. Statistical analyses in **(I-J)** performed using the Wilcoxon matched-pairs signed rank test. See Supplemental Tables for: for IgV_H_-Seq, rmAb and UCA sequences (Table S2), lineage assignments (Table S2), and L-SPR data (Table S4).

### Linking the allospecific tissue and circulating B cells to the systemic alloantibody response

Circulating DSA, which can be produced by terminally differentiated ASCs or by Bmem cells that rapidly differentiate into ASCs upon stimulation, contribute to B lineage-dependent allograft pathology.^2,4^ Since prior vaccination and infection studies have revealed that the BCR repertoires of ASCs and Bmem may differ,^46,47^ we compared the Ig proteome of the systemic plasma A*01:01-specific polyclonal Ab from recipient N006 to the BCR repertoire of the A*01:01-specific B cells and ASCs from kidney and blood of recipient N006. We first affinity-purified the polyclonal A*01:01-specific Abs (appAbs) from plasma of recipient N006 and then enzymatically digested the appAbs to peptides that were subsequently analyzed using liquid chromatography-tandem mass spectrometry (LC-MS/MS). We aligned the A*01:01-specific appAb peptide sequences to the AA sequences encoded by the IgV_H_ nucleotide sequences derived from the A*01:01-specific BCR clonal lineages identified in recipient N006. Using this approach, we identified overlapping A*01:01-specific appAb peptides that matched the sequences of the BCRs expressed by the A*01:01-specific B cells and ASCs found in recipient N006 kidney and blood (Table S5). Importantly, this overlap included matched identical sequences across the hypervariable HCDR3 regions of the rmAbs derived from recipient N006 A*01:01-specific B cells. For example, AA sequences of peptides recovered from the A*01:01-specific appAbs were identical to IgV_H_ AA sequences from two different Lin4 rmAbs (D01 and G22) cloned from blood Bmem (Figure 3A-C). This finding was not limited to Lin4 as we identified many circulating polyclonal appAb peptide sequences that could be mapped to A*01:01-specific rmAbs cloned from Lin14 and Lin6 blood Bmem (Figure 3D,E). In fact, we mapped appAb peptides to IgV_H_ sequences from 4 different A*01:01-specific lineages (Figure 3F) that included kidney- and blood-derived B cells and ASCs. Thus, circulating DSA was clonally related to the ongoing donor-specific blood and kidney B cell and ASC responses.

**Figure 3.**
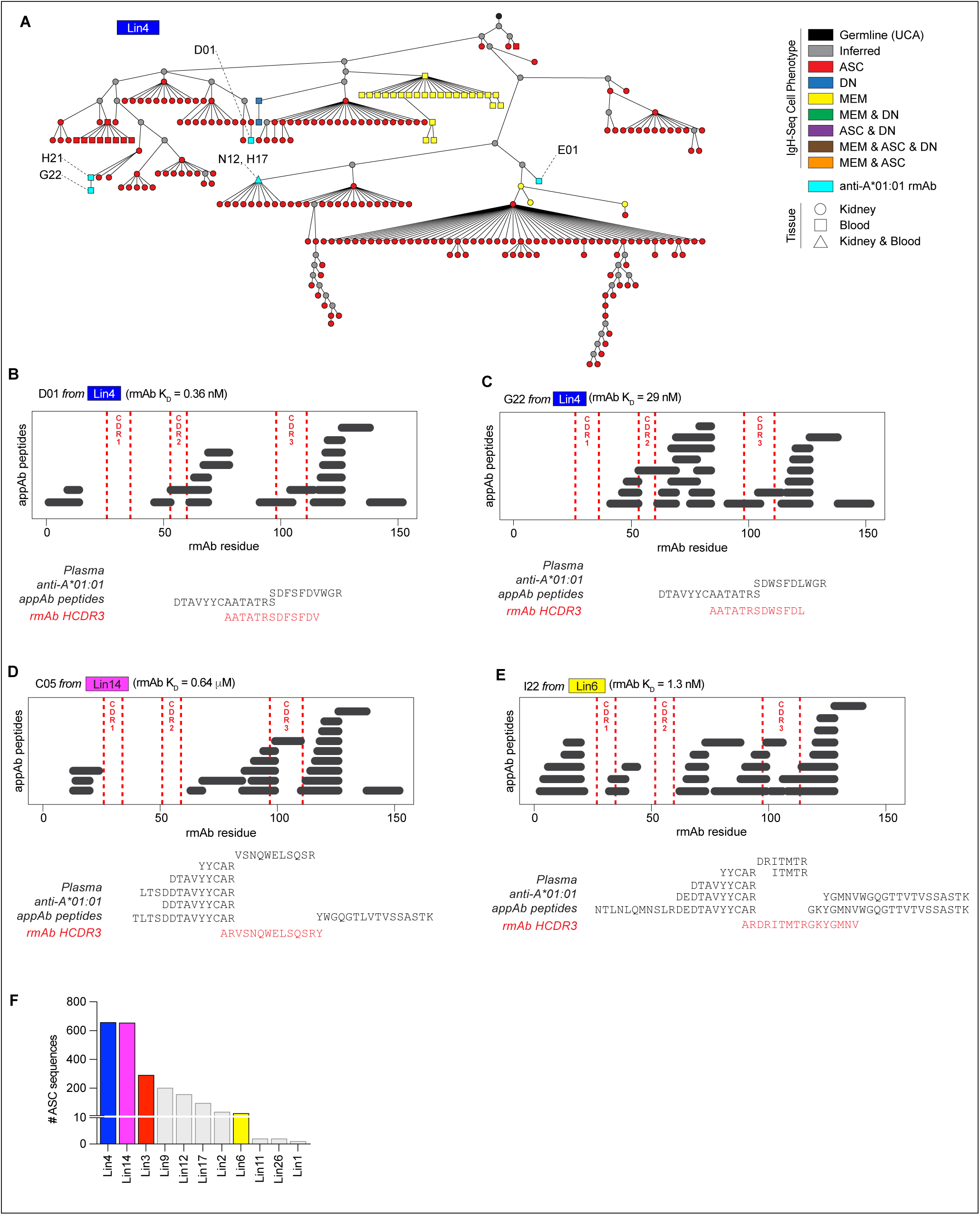
Direct linkage of Donor Specific Abs to ASCs and B cells from kidney and blood. **(A)** Phylogenetic tree showing Lin4 A*01:01-specific B cells and ASCs from kidney and blood. See description of tree symbols and lines in Figure 2A. **(B-E)** Alignment of plasma proteomics data to A*01:01-specific rmAb sequences. Peptide sequences derived from LC-MS/MS analysis of enzymatically digested A*01:01-specific polyclonal appAbs from recipient N006 were aligned against sequences of Lin4 (**B-C**), Lin14 (**D**), and Lin6 (**E**) A*01:01-specific rmAbs cloned from B cells isolated from recipient N006. Exact peptide matches are plotted by position within the rmAb VDJ AA sequence. Peptide sequences overlapping the rmAb HCDR3 shown. L-SPR-calculated K_D_ of the A*01:01-specific rmAbs indicated. **(F)** Column plot showing number of IgV_H_ sequences derived from bulk sorted blood and kidney ASCs in 11 A*01:01-specific clonal lineages. A*01:01-specific lineages containing AA sequences that were also detected in the A*01:01-specific polyclonal appAb proteome are colorized. See Table S5 for proteomics data.

### Dominant HLA reactivity patterns within the allospecific B cell lineages

Our data showed that the B cell and ASC response to the donor HLA-A alloantigen was robust and included at least 14 distinct B cell clonal lineages. These data suggested that the genetically diverse alloreactive B cells might be specific for different epitopes within the mismatched HLA protein. If so, we predicted that the rmAbs derived from distinct clonal lineages would exhibit differing patterns of reactivity for other third party (neither host nor donor) HLA-A proteins that were also mismatched to self-HLA at one or more AA residues. We identified 13 residues that were double mismatched (2MM) between the HLA-A expressed by the donor kidney (A*01:01) and the self-HLA expressed by recipient N006 (A*24:02 / A*30:01). Of the 13 2MM AA residues, 7 AA were solvent-accessible (defined as ≥50 Å^2^ solvent-accessible surface area) and could potentially be targeted by an Ab.^15^ Therefore, we hypothesized that the collection of 50 rmAbs would exhibit different patterns of HLA reactivity that targeted these seven individual mismatched residues. To test this, we examined the binding of the 50 A*01:01-specific rmAbs against a panel of 21 different HLA-A molecules (Figure 4A, Figure S3B, Table S3). Consistent with our hypothesis, we observed that the rmAbs displayed 17 distinct HLA-A reactivity patterns (Figure 4A, Figure S3C). Importantly, when we compared the reactivity profiles of the 50 rmAbs cloned from Bmem and DN isolated from recipient N006 to the reactivity profile of the A*01:01-specific polyclonal appAbs found in circulation of the same individual, we found that the patterns were strikingly similar – with the rmAbs and the appAbs showing the highest reactivity towards three HLA-A proteins: A*01:01, A*36:01 and A*80:01 (Figure 4B). Thus, the HLA reactivity profile of the systemic appAbs was very similar to the aggregated profile of the collection of rmAbs that were cloned from 50 A*01:01-specific blood and kidney B cells.

**Figure 4.**
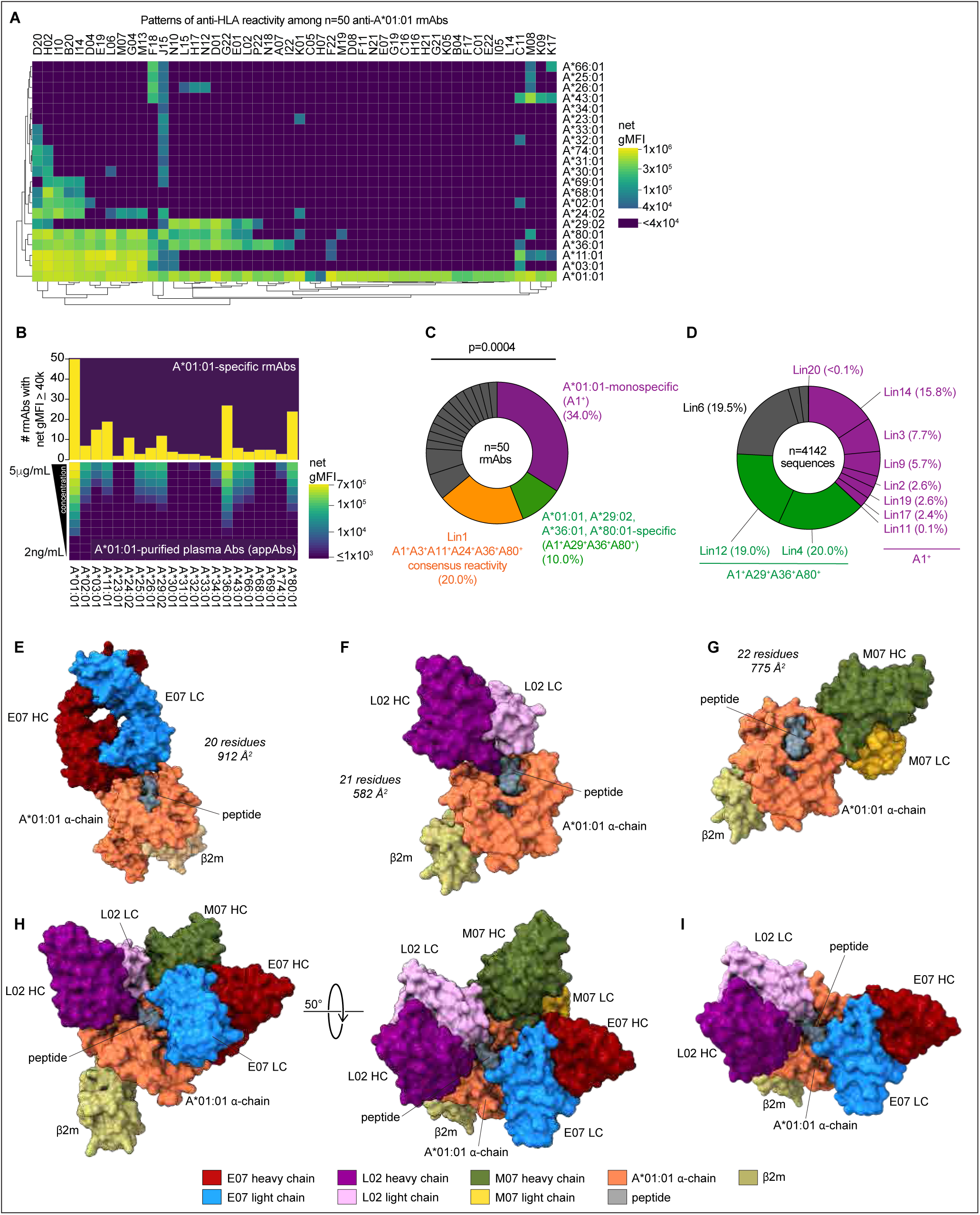
HLA-A*01:01-specific B cells exhibit immunodominant HLA reactivity patterns and recognize unique A*01:01 epitopes. **(A)** HLA-A reactivity profile by 50 A*01:01-specific rmAbs cloned from B cells isolated from recipient N006 KD and blood. Data reported as net gMFI of rmAb binding to microbeads displaying 21 individual HLA-A alleles following normalization against a panel of negative (influenza-specific) control human rmAbs (see *Methods*). Data clustered using Euclidean distance. **(B)** Comparison between HLA reactivity profiles of 50 A*01:01-specific rmAbs and circulating affinity-purified A*01:01-specific appAb isolated from recipient N006 plasma. Upper plot shows number of rmAbs that bound (net gMFI ≥40,000) to each HLA-A protein bead. Bottom plot shows net gMFI of appAb (diluted 2-fold from 5 µg/mL to 2 ng/mL) binding to each HLA-A protein bead. **(C)** Pie chart depicting the most common HLA reactivity patterns for 50 A*01:01-specific rmAbs cloned from recipient N006 kidney and blood B cells. rmAb HLA reactivity patterns were clustered by converting net gMFI of each allele into binary variables (net gMFI ≥40,000 = positive or net gMFI <40,000 = negative). **(D)** Pie chart depicting the most common HLA reactivity patterns for the 14 A*01:01-specific IgV_H_ lineages (n=4142 sequences). For each clonal lineage, a dominant pattern of anti-HLA reactivity (See Figures S3, S4) was assigned by identifying the specific HLA-A proteins bound the rmAbs associated with each clonal lineage. **(E-G)** Structures showing IgH and IgL chains of kidney Bmem cell derived A*01:01-specific rmAbs E07 (**E**), L02 (**F**) and M07 (**G**) bound to HLA-A*01:01. HLA-A*01:01 α-chain (coral), β2-microglobulin (β2m, khaki), and peptide (slate gray). Number of A*01:01 α-chain AA residues contained within each epitope and the amount of A*01:01 α-chain surface area buried upon binding of each rmAb indicated. **(H)** Merged, simulated structure showing binding of rmAbs E07, L02, and M07 to HLA-A*01:01. (**I**) Merged, simulated structure showing binding of rmAbs E07 and L02 to HLA-A*01:01. Statistical analysis in **(C)** tested a null hypothesis that there would be 7 different patterns of reactivity corresponding to the 7 solvent-accessible and double-mismatched residues expressed by donor A*01:01 relative to recipient HLA-A*24:02 and A*30:01. Data analyzed using a binomial test with an expected frequency for an A*01:01-monospecific pattern of 14.3%. The confidence interval of the proportion was calculated using the Wilson/Brown method. An A*01:01 monospecific pattern of reactivity was observed significantly more often than expected (34.0% vs. 14.3%, p=0.0004). See Supplemental Figures and Tables for: HLA reactivity patterns (Figure S3, S4) cytometric bead data (Table S3, Figure S3, Figure S4); sequencing data (Table S2); and structural data (Table S6).

Although we observed multiple reactivity patterns among the A*01:01-specific rmAbs (Figure 4A,B, Figure S3C), some patterns were more highly represented within the rmAb collection. For example, the A*01:01-monospecific reactivity pattern (referred to as A1^+^) included 17/50 (34.0%) rmAbs that bound A*01:01 alone (Figure 4C). The 17 rmAbs exhibiting an A1^+^ monospecific reactivity pattern were derived from eight distinct clonal lineages (Figure S4A). Similarly, 5/50 (10.0%) rmAbs spread across two distinct clonal lineages (Lin4 and Lin12) exhibited an A*01:01^+^A*29:02^+^A*36:01^+^A*80:01^+^ (A1^+^A29^+^A36^+^A80^+^) reactivity pattern (Figure 4C, Figure S4B). Finally, a third lineage (Lin1), which contained ten clonally related rmAbs, exhibited consensus binding to six HLA-A alleles: A*01:01^+^A*03:01^+^A*11:01^+^A*24:02^+^A*36:01^+^A*80:01^+^ (Figure 4C, Figure S4C).

Next, we investigated whether specific HLA reactivity patterns were associated with the largest lineages in the A*01:01-specific IgV_H_-seq database. Lin4, which represented 20.0% of all IgV_H_ sequences in the A*01:01 specific IgV_H_-seq database and contained four rmAbs (Figure 4D), displayed A1^+^A29^+^A36^+^A80^+^ reactivity (Figure S4B). Lin12, which also displayed A1^+^A29^+^A36^+^A80^+^ reactivity (Figure S4B), contained one rmAb and comprised 19.0% of A*01:01-specific IgV_H_ sequences (Figure 4D). Thus, five A1^+^A29^+^A36^+^A80^+^ rmAbs were linked to two clonal lineages that collectively constituted 39.0% (Figure 4D) of all A*01:01-specific IgV_H_ sequences. Similarly, the eight clonal lineages containing the 17 A1^+^ monospecific rmAbs (Figures S4A) represented 36.9% of the entire A*01:01-specific IgV_H_ repertoire (Figure 4D). Thus, despite appreciable diversity in the HLA-reactivity patterns within the A*01:01-specific B cell response, two patterns of HLA reactivity, which together encompassed 75.9% of all A*01:01-specific IgV_H_ sequences in our database, dominated the A*01:01-specific IgV_H_-Seq repertoire. These results suggested that the B cell and Ab response to the mismatched HLA-A protein might be focused on a limited number of immunodominant HLA-A epitopes.

### HLA epitope recognition by immunodominant allospecific B cells

Although mismatched HLA residues are necessary to generate alloimmunity^21^ and the number of HLA residue mismatches between self and non-self positively correlates with the formation of DSA responses,^22^ the relative contributions and immunogenicity of individual mismatched residues remain unknown.^23–25^ Our clonal lineage and HLA reactivity data suggested strongly that the B cell alloresponse might be focused on a relatively small number of highly “immunogenic” mismatched residues. To test this idea, we used X-ray crystallography and cryo-electron microscopy (cryoEM) to determine the structures of 3 representative rmAb/HLA-A*01:01 complexes. These included the Lin2 E07 rmAb (Figure 4E), which exhibited the immunodominant A1^+^ monospecific pattern of reactivity (Figure S4A) and was derived from a kidney A*01:01-specific Bmem, and the Lin12 L02 rmAb (Figure 4F), which exhibited the immunodominant A1^+^A29^+^A36^+^A80^+^ pattern of reactivity (Figure S4B) and was also derived from a kidney Bmem. Finally, we examined the kidney Bmem-derived rmAb M07 (Figure 4G), which was part of Lin1 that included 20.0% (10/50) of the rmAb collection and exhibited the A1^+^A3^+^A11^+^A24^+^A36^+^A80^+^ reactivity profile (Figure S4C).

To characterize the structural epitope(s) in HLA-A*01:01, we defined the epitope as residues with an interatomic distance of ≤5 Å between a heavy atom in the Fab and a heavy atom in the HLA complex or any residues in the HLA complex with solvent-accessible surface area that was buried in whole or in part in the interface between the Fab and HLA-A*01:01 (Table S6). Using these criteria, we found that the footprints of the HLA epitopes bound by the three rmAbs were consistent with the size of typical protein epitopes^15^ with the rmAbs burying 912 Å^2^ (E07), 582 Å^2^ (L02), and 775 Å^2^ (M07) of solvent accessible surface area within the HLA-A*01:01 α-chain (Figure 4E-G, Table S6). The number of AA residues (n=20-22 A*01:01 α-chain residues) present in each of the HLA epitopes (Figure 4E-G) was equivalent to other previously described protein epitopes.^15,48^ Consistent with the different HLA reactivity patterns exhibited by each rmAb, we found that the rmAbs bound three distinct epitopes on A*01:01, with little overlap in the binding footprints (Figure 4H). For example, the HLA-A*01:01 epitopes recognized by rmAbs E07 (Lin2, A1^+^ monoreactivity pattern) and M07 (Lin 1, A1^+^A3^+^A11^+^A24^+^A36^+^A80^+^ reactivity pattern) only overlapped by four AAs (D129, R131, E154, E157). By contrast, rmAb L02 (Lin12, A1^+^A29^+^A36^+^A80^+^ reactivity pattern) and rmAb E07 (A1^+^ binding pattern) showed no overlap in their epitope footprints (Figure 4I). Thus, the three kidney Bmem derived rmAbs, which were representative of the three most dominant reactivity profiles, recognized unique, largely non-overlapping epitopes on A*01:01.

### Immune focusing of the alloreactive B cell response on the crown of HLA-A

Our data showed that the three rmAbs, which recognized largely non-overlapping epitopes on the top or crown of HLA A*01:01, exhibited three distinct patterns of reactivity toward third party HLA-A alleles (Fig. 5A-C, Table S3). Since polymorphic residues in antigens may contribute to immune focusing and immunodominant B cell and ASC responses,^19,20^ we predicted that binding of the rmAbs to A*01:01 and other third-party HLA-A proteins would be dictated by the presence of one or more mismatched residues that were present in the A*01:01 epitope recognized by the rmAb and shared by the mismatched HLA-A (A*01:01) plus the other third party HLA-A proteins that were recognized by the same rmAb. To define the mismatched residues, we compared A*01:01 (donor) residues to A*24:02 and A*30:01 (self) residues expressed by recipient N006 and identified 2MM residues that differed from both self-alleles as well as 1MM residues that differed between donor HLA and one of the self-HLA alleles. In agreement with our hypothesis, we identified multiple 2MM and 1MM residues that could explain the binding patterns of each of the 3 rmAbs (Figure 5D-I). For example, the A1^+^ monospecific binding pattern of E07 (Figure 5A) could be potentially explained by three solvent accessible residues (Figure 5D), V158, R163 (both 2MM residues) and D166 (1MM residue) that were present in the A*01:01 epitope recognized by E07, were buried by or formed H-bonds with the bound E07 (R163, D166; Figure 5E), and were not co-expressed by any of the other tested third-party class I HLA-A alleles (Figure 5D, Table S3). Thus, the data suggested that both 2MM and 1MM residues within the A*01:01 epitope recognized by E07 had the potential to influence the specificity and reactivity profile of the A*01:01-specific rmAbs. Importantly, these findings were not limited to the A*01:01/E07 complex as similar results were observed when we examined the epitopic residues and HLA binding patterns of A*01:01-specific L02 (Figure 5F-G) and M07 (Figure 5H-I) rmAbs.

**Figure 5.**
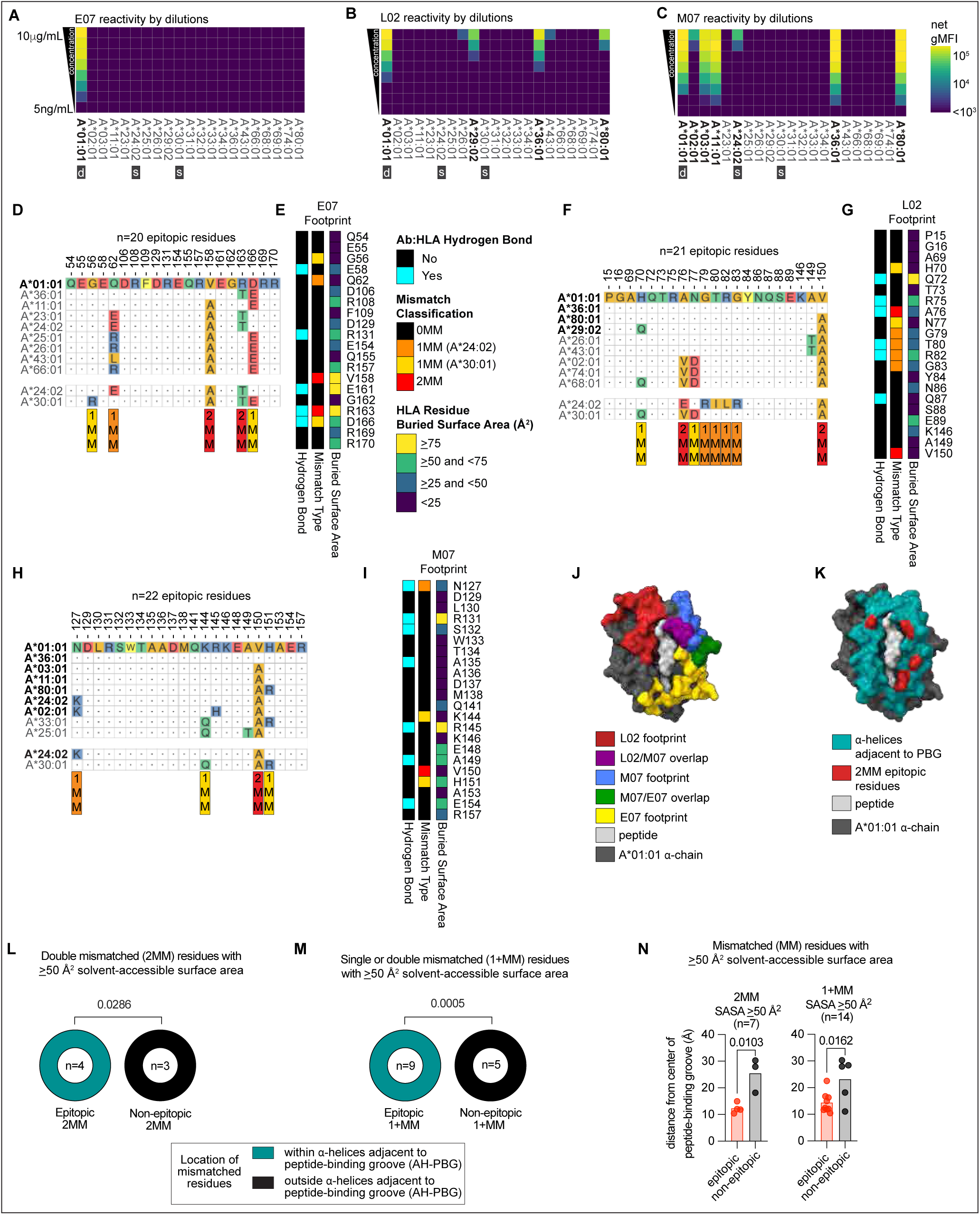
Immune focusing of A*01:01 specific B cells on the crown of HLA-A*01:01. **(A-C)** HLA-A reactivity profiles showing binding of kidney Bmem cell derived rmAbs E07 (**A**), L02 (**B**) and M07 (**C**) to microbeads displaying 21 individual HLA-A proteins. Data reported as net gMFI of the average of 2 technical replicates of rmAbs that were diluted from 10 µg/mL to 5 ng/mL. Donor allele (“d”) and self-alleles (“s”) indicated. The HLA reactivity profile for each rmAb is defined as all HLA-A alleles (bolded) with a net gMFI ≥40,000 following incubation with the rmAbs at 10 µg/mL. **(D-I)** Characterization of HLA-A*01:01 epitopic AA residues bound by rmAbs E07, L02 and M07 derived from recipient N006 kidney Bmem cells. Panels **D**, **F**, **H** highlight the A*01:01 α-chain AA residues (top row, residues numbered 1-274) in each structurally defined HLA-A*01:01 epitope bound by rmAb E07 (**D**, n=20 residues), L02 (**F**, n=21 residues) and M07 (**H**, n=22 residues). The epitopic HLA-A*01:01 residues are aligned with other HLA-A alleles, including self-alleles A*24:02 and A*30:01 and third-party HLA alleles also bound by E07, L02 or M07 (see panels **A-C**). AA are colored based on chemical properties (yellow=aromatic, red=acidic, blue=basic, aliphatic=orange, polar=green). Identical residues between the A*01:01 donor allele and both self-alleles (0 mismatches or 0MM are shown as dots (”.”). Residues mismatched between self, donor or third-party alleles are indicated with the one letter AA code. Residues double-mismatched against both self-alleles (2MM, red) or single-mismatched (1MM) against one self-allele (A*24:02, orange or A*30:01, yellow) indicated. HLA alleles bound by the rmAb (gMFI ≥40,000 at 10 µg/mL) are bolded. Heatmaps in panels **E** (rmAb E07), **G** (rmAb L02) and **I** (rmAb M07) depict the A*01:01 epitopic residues involved in hydrogen bonds with the rmAbs (left heatmap), the 0MM, 1MM or 2MM A*01:01 epitopic residues (middle heatmap) and the amount of surface area of each epitopic residue that is buried by rmAb binding (right heatmap). **(J)** Depiction of the three epitopic footprints of E07, L02, and M07 bound to A*01:01. Regions of overlap between epitopes indicated. **(K)** Location of the four 2MM residues (A76, V150, V158, R163) within the A*01:01 epitopes bound by rmAbs E07, L02, or M07. Peptide (grey), 2MM residues (red), AH-PBG region (teal). **(L,M)** Pie charts depicting the location of 2MM and 1MM plus 2MM (1+MM) residues within the HLA epitopes bound by E07, L02, and M07, comparing epitopic MMs (left pie chart) versus non-epitopic MMs (right pie chart). Each pie chart indicates the proportion of MMs contained within the AH-PBG region (teal) or contained outside the AH-PBG region (black). **(L)** depicts 2MM residues with ≥50 Å^2^ of solvent-accessible surface area (n=7). **(M)** depicts all 1MM and 2MM residues (1+MM) with ≥50Å^2^ of solvent-accessible surface area (n=14). **(N)** Distance of solvent-accessible 2MM residues (left, n=7) or 1+MM residues (right, n=14) from the center of the PBG in A*01:01 (see Figure S3A for all 2MM residues). Residues contained in the HLA epitopes bound by E07, L02, or M07 (red) and residues outside of the HLA epitopes recognized by these 3 rmAbs (black). Statistical analysis **(L,M)** performed using Fisher’s exact test. Comparisons in **(N)** used an unpaired t-test. All distributions passed normality (Shapiro-Wilk test). See Supporting Figures and Tables for: A*01:01-specific rmAb cytometric bead array data (Table S3) and additional A*01:01 epitope structure data (Figure S5 and Table S6).

We next asked whether conventional, non-machine learning-based epitope prediction tools would accurately identify the solvent-accessible 1MM and 2MM residues that we knew were contained within the structurally defined HLA epitopes recognized by the three rmAbs. Interestingly, common approaches used to predict HLA residue immunogenicity,^49–51^ including protrusion index, hydrophilicity, isoelectric point, and structural similarity, failed to discriminate (Figure S5A,B) between the mismatched residues that were encompassed within the epitopes (epitopic) recognized by rmAb relative to the mismatched residues that were located outside of the A*01:01 epitopes (non-epitopic). Since the standard immunogenicity tools did not accurately predict which mismatched residues were contained within the structurally defined HLA/alloAb epitope, we asked whether the topography of HLA might influence immunogenicity and immune focusing to certain subregions of the alloantigen as has been reported for other antigen/Ab complexes.^52^ We therefore assessed the position of the three epitopes recognized by the three rmAbs within the quaternary structure of HLA-A*01:01, focusing on the mismatched solvent accessible residues (defined as residues with ≥50 Å^2^ exposed solvent accessible surface area) that could form close interactions with Ab molecules. We found that all three rmAbs made multiple contacts (Figure 5J) with the HLA-A α-helices (α1 domain residues 50-85 and α2 domain residues 138-175) that frame the peptide-binding groove (PBG) and form the membrane-distal “crown” of HLA (AH-PBG region Figure S3A).^53^ We further observed that solvent-accessible residue mismatches recognized by E07, L02 and M07 (i.e., epitopic mismatches) were significantly more likely to reside within the AH-PBG on the crown of the HLA molecule as compared to mismatches that were not contained within the HLA epitopes bound by these Abs. This was true for the four solvent-accessible 2MM epitopic residues (A76 [L02], V150 [L02 and M07], V158 [E07], and R163 [E07]) (Figure 5K) located within the AH-PBG region (p=0.0286, Figure 5L) as well for the 9 epitopic 1MM and 2MM residues (p=0.005, Figure 5M). Consistent with this observation, solvent-accessible 2MM residues targeted by these rmAbs were also significantly closer to the center of the PBG (p=0.0103). This was still true when we included both 1MM and 2MM residues (p=0.0162) (Figure 5N, Figure S5C,D) and when we applied a lower solvent accessibility surface area threshold^54^ of 30 Å^2^ (Figure S5E-G). Thus, these data suggest that the immunogenicity of specific 1MM and 2MM A*01:01 residues may be controlled by a combination of HLA topography and residue solvent accessibility.

### AlloHLA-specific B cells target common immunodominant epitopes on A*01:01

Our data indicated that the mismatched solvent-accessible A*01:01 residues encompassed within the epitopes recognized by the three representative rmAbs were located on the crown of the HLA-A molecule within the AH-PBG region. We predicted that the other rmAbs that were part of the same clonal lineages as E07, L02 or M07 would recognize the same “immunodominant” sites on the HLA-A α-chain. In addition, we anticipated that rmAbs that were derived from clonal lineages unrelated to E07, L02 or M07, but exhibited the same HLA-A third-party binding patterns as our representative rmAbs, should also recognize the same immunodominant epitope(s). We tested these postulates with two of the A*01:01-specific rmAbs, E07 and L02, which were derived from distinct clonal lineages (Figure 2A,C), exhibited different HLA reactivity patterns (Figure 5A,B), and recognized completely non-overlapping epitopes on A*01:01 (Figure 4I). We first generated chimeric (c) rmAb versions of E07 and L02 by fusing the IgV_H_ and IgV_L_ domains from the human E07 or L02 rmAbs to the mouse IgG1 constant region (cE07 or cL02, see *Methods*). We next preincubated A*01:01-coated microbeads with one or both chimeric rmAbs and then incubated the beads with one of the 22 test rmAbs, which included 17 A1^+^-monospecific human rmAbs from 8 distinct clonal lineages (Figure S4A) and the 5 A1^+^A29^+^A36^+^A80^+^ reactive human rmAbs from Lin4 and Lin12 (Figure S4B). To measure whether the presence of the chimeric blocking Abs impeded binding by the test human rmAb, we used fluorochrome-labeled secondary Abs specific for the constant region of *human* IgG, which selectively recognized the human A*01:01-specific “test” rmAbs. For a negative control blocking Ab, we used a murine anti-human HLA Ab (mouse IgG2a clone W6/32)^55^ that recognizes the β2m residues 3 and 89 and residue 121 in the α2 domain of the HLA α-chain (which is located outside of the AH-PBG).^56,57^ As expected, W6/32 effectively bound to HLA-A*01:01 (Figure S6A) but only minimally inhibited the binding of any of the A*01:01-specific human rmAbs – with a median inhibition of only 2% (Figure 6A). Also, as expected, the cE07 rmAb completely inhibited the binding of its non-chimeric “parent” human Lin2-derived E07 rmAb (Figure 6A). In addition, cE07 rmAb also blocked the binding of the additional 6 Lin2-derived rmAbs (Figure 6A), indicating that all rmAbs in this clonal lineage likely recognized the same epitope. Moreover, cE07 rmAb almost completely inhibited (>90%) binding of 9/10 of the other A1^+^ monospecific rmAbs (Figure 6A,B), which were derived from an additional seven clonal lineages (Figure S4A), thereby suggesting that these genetically unrelated A1^+^ monospecific rmAbs also likely recognized the same or a closely overlapping epitope as the A1^+^ monospecific E07 rmAb. Consistent with our structural data showing that E07 and L02 recognize distinct, non-overlapping A*01:01 epitopes (Figure 4I), cE07 rmAb did not greatly inhibit the binding of human Lin12 L02 rmAb or 4 other A1^+^A29^+^A36^+^A80^+^-specific rmAbs that were part of the separate Lin4 genetic lineage (Figure 6A,B). By contrast, cL02 (an A1^+^A29^+^A36^+^A80^+^-specific rmAb) inhibited the binding of its non-chimeric “parent” Lin12 human L02 rmAb (Figure 6A). The cL02 rmAb also blocked binding of the four A1^+^A29^+^A36^+^A80^+^-specific rmAbs from Lin4 (Figure 6A,C). By contrast, cL02 inhibited the binding of the A1^+^ monospecific rmAbs significantly (p<0.0001) less well (Figure 6C). Therefore, these data show that rmAbs that are either clonally related or share identical patterns of reactivity seem to recognize overlapping or neighboring epitopes on A*01:01.

**Figure 6.**
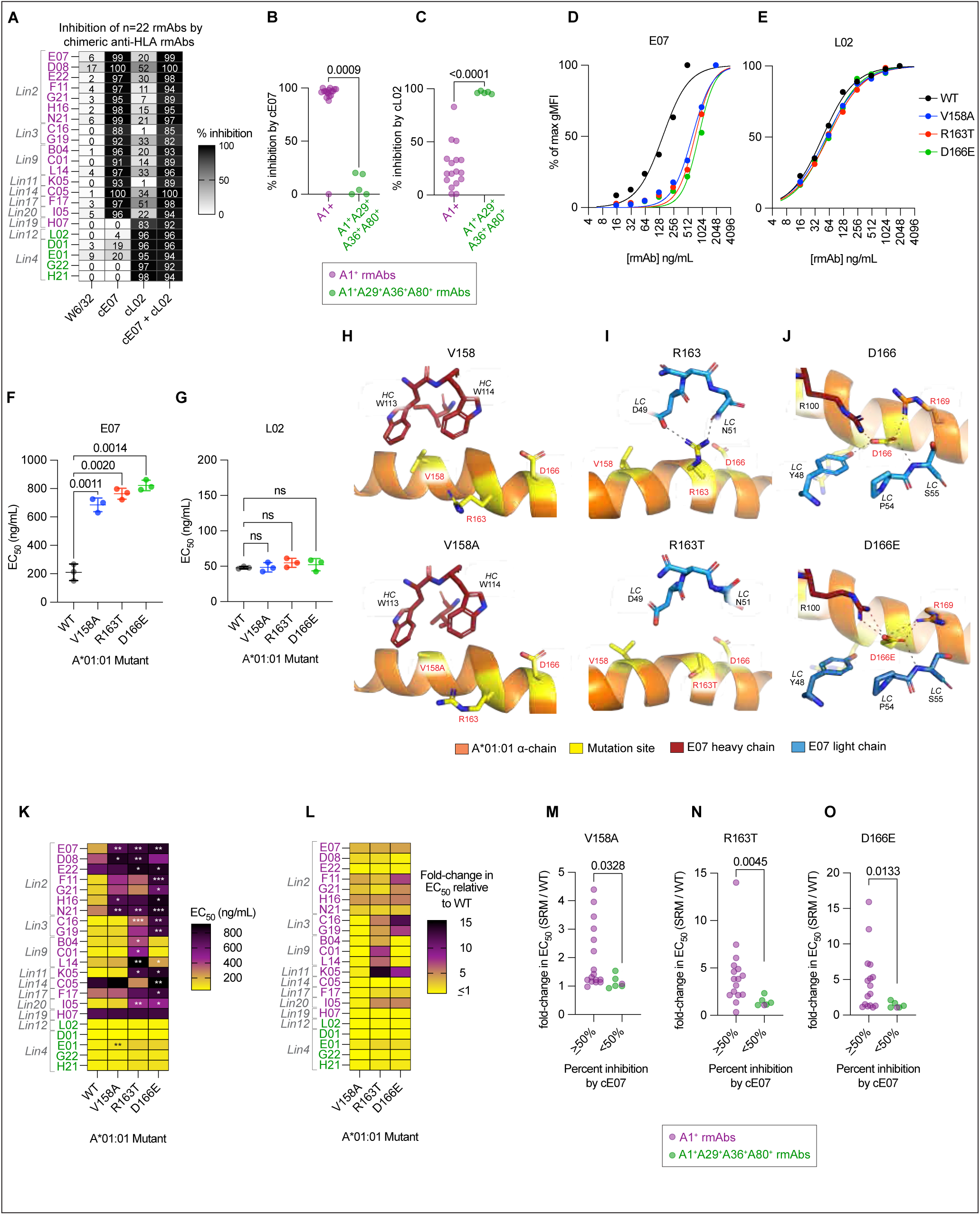
Convergent recognition of HLA-A*01:01 crown-localized epitopes by multiple distinct B cell clonal lineages. **(A)** Chimeric mouse/human rmAbs cE07 and cL02 block binding of A*01:01 specific rmAbs derived from 10 independent B cell clonal lineages. HLA-A*01:01 microbeads were blocked with chimeric A*01:01-specific cE07 and/or cL02 rmAbs, then stained with human A*01:01-specific test human rmAbs (n=22, Figure S4A-B) that were subsequently detected with fluorochrome-labeled anti-human IgG. Data reported as percent inhibition of test rmAb binding (mean of 22 technical replicates) and are representative of 2 independent experiments. Test human A*01:01-specific rmAbs subdivided based on clonal lineage and HLA reactivity patterns. W6/32, a murine Ab specific for human β2m, included as a negative control for inhibition. **(B-C)** Comparison between HLA binding reactivity pattern of test rmAbs and inhibition of test rmAb binding to A*01:01 mediated by chimeric rmAbs cE07 or cL02. Data shown as % inhibition with test rmAbs grouped by HLA binding patterns. **(D-G)** Binding to HLA-A*01:01 wild-type (WT) protein and single residue mutants of A*01:01 (V158A, R163T, D166E) by rmAbs E07 (**D**) and L02 (**E**). Data shown as increasing concentrations of the rmAbs (x-axis) and percent of maximum gMFI for binding to A*01:01 (y-axis). Binding curves were fit to each rmAb:protein combination and used for EC_50_ calculations ((**F-G**), see *Methods* for details). Data shown for three independent experiments for each rmAb:protein combination, with lines indicating mean with standard deviation. **(H-J)** Interactions of rmAb E07 with 2MM and 1MM residues in A*01:01. Top cartoons show interactions between residues in the E07 IgV_H_ domain (heavy chain [HC], firebrick red) or IgV_L_ domain (light chain [LC], blue) and A*01:01 (orange cartoon) with selected mismatched AA residues in A*01:01 (yellow cartoon with sticks), including 2MM V158 (**H**), 2MM R163 (**I**), and 1MM D166 (**J**). Bottom cartoons show reduced contact between E07 and HLA structures harboring A*24:02/A*30:01 self-reversion mutations V158A (**H**), R163T (**I**), or D166E (**J**). Polar interactions noted by dashed lines. (**K**) Heatmap showing EC_50_ values for 22 test rmAbs binding to WT and mutant A*01:01 microbeads. Data shown as mean of three independent experiments for each rmAb:protein combination. Test rmAbs grouped by clonal lineage and HLA reactivity pattern. (**L**) Heatmap showing relative reduction in binding to A*01:01 mutants by 22 A*01:01 specific rmAbs. Data reported as the fold change in EC_50_ values (mutant EC_50_/WT EC_50_) for each rmAb and shown as mean of three independent experiments for each rmAb:protein combination. Test rmAbs grouped by clonal lineage and HLA reactivity pattern. **(M-O)** Comparison of test rmAb binding to A*01:01 mutants to chimeric cE07 rmAb mediated inhibition of test rmAb binding to A*01:01. Binding of test rmAbs to A*01:01 mutants shown as fold-change in EC_50_ values (mutant EC_50_/WT EC_50_). Test rmAbs grouped based upon their percent inhibition (250% or <50%) by cE07. Test rmAbs colored based upon anti-HLA binding patterns. Statistical analysis (**B-C, M-O**) performed using a Mann-Whitney test. Nonlinear regression with a sigmoidal four-parameter logistic curve (**D-G,K**) used to fit binding for each rmAb:protein combination and calculate EC_50_ values. A*01:01 mutant EC_50_ values were compared against the WT EC_50_ values with Brown-Forsythe and Welch ANOVA tests (**F,G,K**) with Dunnett’s T3 multiple comparisons test. In **(K)**, statistical significance is indicated with asterisks: * P≤0.05, ** P<0.01, *** P<0.001. See Supporting Figures for: A*01:01-specific rmAb cytometric bead array data (Figure S4) and additional rmAb binding data and A*01:01 epitope structure data (Figure S6).

While our data suggested that B cells derived from distinct clonal lineages converged on the same limited set of immunodominant HLA epitopes, we could not rule out the possibility that binding of a chimeric rmAb simply sterically hindered the binding of other rmAbs to non-overlapping epitopes. To test this, we first identified the five 1MM and 2MM HLA-A*01:01 residues that were present in the E07 structurally defined epitope (Figure 5D) and were not present within the L02 structurally defined epitope (Figure 5F). From those five mismatched residues, we focused on the 2MM residues V158, R163 and the 1MM residue D166as V158, R163 and D166 formed the closest contacts with E07, had the largest surface areas buried upon E07 binding, were located within the AH-PBG region, and formed H-bonds (R163 and D166) with E07 (Figure 5E, Figure S6B). We then generated three recombinant single residue mutants of A*01:01 by reverting V158, R163 and D166 to a “self” HLA-A AA residue expressed by recipient N006 at each site (V158A, R163T, D166E). For example, the 2MM V158 was changed from the A*01:01-encoded valine to the alanine that is expressed by both self-HLA (A*24:02 and A*30:01) expressed by recipient N006. Likewise, the 1MM D166 was reverted from the donor A*01:01 aspartate to the glutamate that is expressed by recipient N006 self-HLA allele, A*30:01.

We next tested the binding activity of E07 and L02 against these A*01:01 mutants by calculating the EC_50_ value for each rmAb against the WT (non-mutated wild-type) A*01:01 and the V158A, R163T, and D166E A*01:01 mutants (Figure 6D-G). Consistent with the fact that all three mutant HLA featured single AA reversions within the epitope recognized by E07, E07 showed significantly decreased binding against all three A*01:01 mutants relative to the WT A*01:01 (Figure 6D,F). By contrast, L02 showed equivalent binding against the A*01:01 mutants relative to WT A*01:01 (Figure 6E,G). These results fit well with structural simulation data showing that the single point mutations in A*01:01 disrupted hydrophobic interactions between A*01:01 and the E07 H-chain (V158A, Figure 6H) and hydrogen bonds between A*01:01 and the E07 L-chain (R163T and D166E, Figure 6I,J).To confirm these results, we used L-SPR to measure the K_D_ of E07 and L02 for binding to WT and mutant A*01:01. We observed that the affinity of E07 binding to each A*01:01 mutants was approximately 20-fold lower when compared to E07 binding to WT A*01:01 (Figure S6C) while the affinity of L02 binding to A*01:01 WT was similar to that observed for binding to the A*01:01 mutants (Figure S6D). Thus, individual mismatched A*01:01 epitopic residues, which are bound by E07 and located in the AH-PBG region, are important for A*01:01 recognition by E07 but not L02.

Given that we now had single residue mutations in A*01:01 that disrupted E07 but not L02 binding to A*01:01, we asked whether the V158A, R163T, and D166E A*01:01 mutations affected binding of the 17 A1^+^ monoreactive rmAbs, which were derived from 8 distinct clonal lineages (Figure 6K,L). We found 16 of these 17 A1^+^-monospecific rmAbs showed significantly decreased binding to at least one of the A*01:01 mutants (Figure 6K) with EC_50_ values that were up to 15-fold higher against the mutant HLAs compared to the WT A*01:01 (Figure 6L). By contrast, 4 of the 5 A1^+^A29^+^A36^+^A80^+^-specific rmAbs showed no decrease in binding to the A*01:01 mutants compared to the WT A*01:01 (Figure 6K). To assess whether decreased binding of the rmAbs to the A*01:01 mutants correlated with cE07-mediated inhibition of binding to A*01:01, we classified the 22 rmAbs by the degree of inhibition by cE07 and the fold-change in the EC_50_ of binding to A*01:01 mutants relative to WT A*01:01. Strikingly, we observed that rmAbs that were most inhibited by cE07 rmAb (≥50% inhibition) also exhibited higher EC_50_ concentrations for binding to the A*01:01 mutants (Figure 6M-O). Thus, >90% of the A1^+^ monospecific rmAbs recognized an epitope in A*01:01 that overlapped with the E07 epitope and that appeared to contain at least one of the mismatched AH-PBG-localized V158, R163, or D166 residues. These data therefore suggest that multiple genetically unrelated clones of B cells converge upon common immunodominant A*01:01 epitopes containing solvent-accessible, exposed residues localized within the AH-PBG region.

### Alloantibody and B cell responses converge on immunodominant HLA epitopes

Our data (Figure 4) showed substantial connectivity between the mismatched allograft A*01:01-specific kidney and blood B cell response and the circulating A*01:01 DSA present in recipient N006. Given that we observed immune focusing of the B cell response on specific solvent-accessible HLA-A residues within the AH-PBG region, we predicted that the circulating DSA response in recipient N006 should also be focused on these residues. To test this hypothesis, we assessed whether the chimeric cE07 and cL02 rmAbs, which were originally cloned from the Bmem found in the kidney allograft of recipient N006, inhibited A*01:01 binding by recipient N006 plasma-derived polyclonal Abs (Figure 7A) or N006’s plasma-derived A*01:01-affinity-purified polyclonal Abs (appAbs) (Figure 7B). We observed that pre-incubation of A*01:01 beads with either cE07 or cL02 partially inhibited A*01:01 binding by the total polyclonal Abs as well as the appAbs (Figure 7A-C). Moreover, pre-incubation of A*01:01 beads with both cE07 and cL02 completely inhibited (94-99%) A*01:01 binding by the total polyclonal Abs and appAbs isolated from recipient N006 plasma (Figure 7C). By contrast, the W6/32 Ab, which does not bind within the AH-PBG region, had a minimal effect on the binding of the systemic polyclonal Abs to A*01:01 (Figure 7C). To confirm that inhibition by cE07 was not solely due to steric hindrance, we next tested whether the A*01:01-specific appAbs purified from the plasma of recipient N006 bound to the A*01:01 single residue mutants V158A, R163T and D166E. We observed a 50-80% reduction in binding by the A*01:01 specific appAbs to the A*01:01 mutants compared to WT A*01:01 (Figure 7D,E). Thus, the alloreactive B cell and serologic responses made by recipient N006 appeared to focus on a small number of immunodominant A*01:01 epitopes, which contained solvent-accessible, mismatched residues within the AH-PBG region.

**Figure 7.**
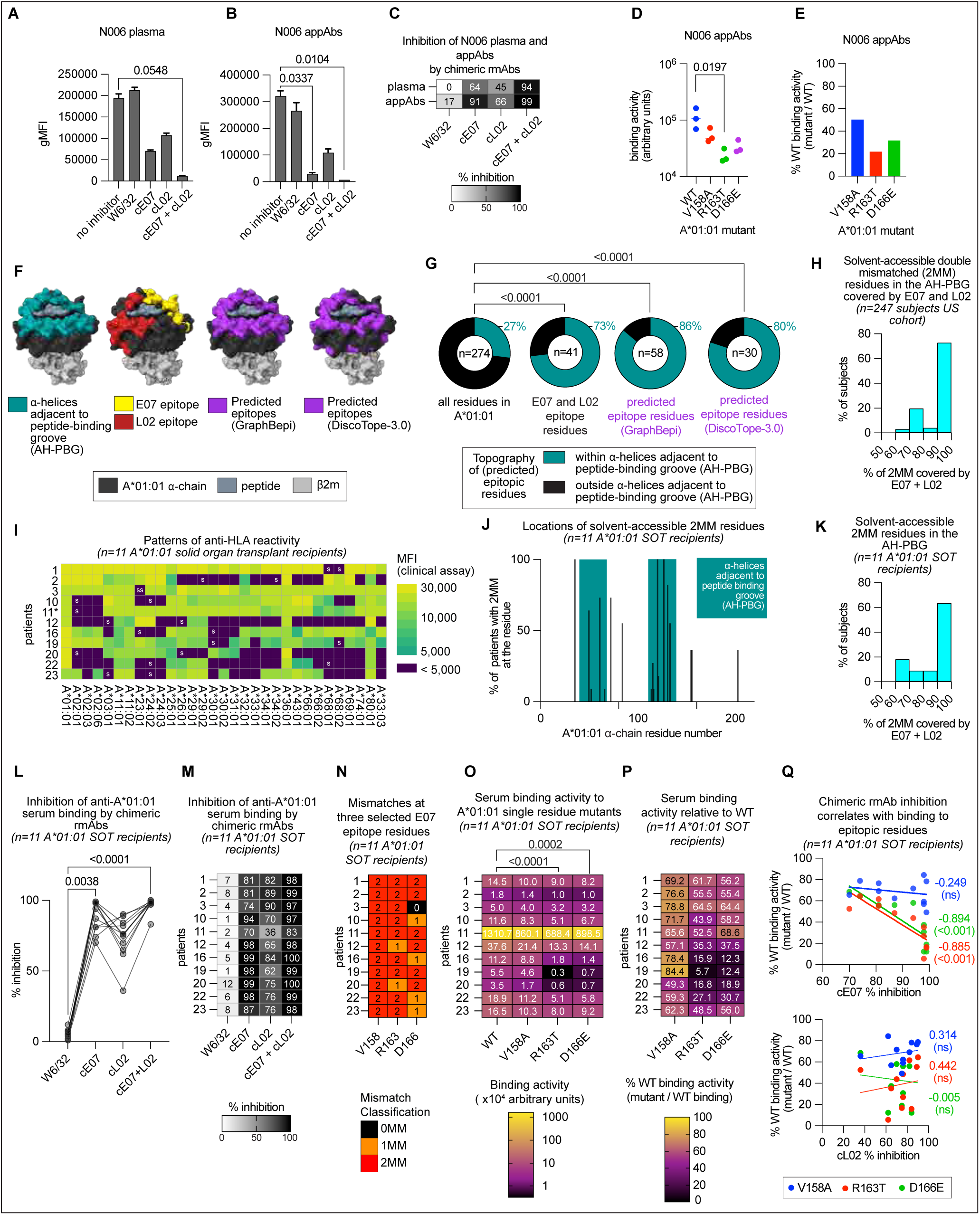
HLA topography enforces convergent immunodominant A*01:01 specific humoral immune responses in a cohort of A*01:01 mismatched transplant recipients. **(A-C)** Chimeric mouse/human rmAbs cE07 and cL02 block binding of circulating polyclonal Abs from recipient N006. Data reported as gMFI (mean plus standard deviation for 22 technical replicates per condition) of binding by total plasma polyclonal Abs (**A**) or affinity-purified A*01:01 binding appAbs (**B**) to A*01:01 coated microbeads. Data in (**C**) reported as percent inhibition of binding and represent the mean of at least two technical replicates. W6/32, a murine Ab specific for human class I HLA, included as a negative control for inhibition. **(D-E)** Binding to WT A*01:01 and mutant A*01:01 (V158A, R163T, D166E) by appAbs from recipient N006. Data are shown as binding activity (arbitrary units, y-axis) for each mutant HLA-A in (**D**) and the percentage of WT binding activity in (**E**). Binding activity was calculated by interpolating binding activity using a standard curve of W6/32 binding for each mutant (Figure S6A). Data representative of three independent experiments for each appAb:protein combination. **(F)** HLA residues contained within the structurally-defined HLA epitopes recognized by E07 (yellow) and L02 (firebrick red) and within the HLA epitopes predicted by GraphBepi and DiscopTope-3.0 (purple) displayed on the HLA-A*01:01 complex with α-chain (black), β2m (light gray), and peptide (dark gray) residues. **(G)** Pie charts comparing structurally defined and computationally predicted HLA-A*01:01 epitopic residues within the 74 AA A*01:01 AH-PBG region. Chart 1 shows proportion of residues located within AH-PBG relative to the total number (n=274) of HLA-A*01:01 α-chain residues. Chart 2 shows proportion of residues within the epitopes recognized by E07 and L02 (n=41) that are localized within the AH-PBG region. Charts 3 and 4 show proportion of residues within the GraphBepi predicted epitope (n=58, pie chart 3), or the DiscoTope-3.0 predicted epitope (n=30, chart 4) that are localized within the AH-PBG region. **(H)** Histogram showing frequency distribution of solvent-accessible (250 Å^2^) 2MM residues that are localized within the A*01:01 epitopes recognized by E07 and L02 rmAbs in a representative cohort (n=247) of HLA-A genotyped subjects who do not express HLA-A*01:01 as a self-allele. **(I)** Heatmap depicting the HLA-A reactivity pattern of serum from 11 transplant recipients who developed a DSA response to A*01:01 expressed by the mismatched donor allograft. Clinical HLA laboratory cytometric bead array data reported as the MFI for serum binding to 31 HLA-A alleles. Self-alleles for each subject indicated (“s”) and subjects homozygous for a given allele indicated with “ss.” For a single subject (recipient 11), one of the self-alleles (A*66:03) was not included in the HLA cytometric bead array. **(J)** Localization of solvent-accessible (250 Å^2^) 2MM A*01:01 residues identified in 11 DSA^+^ transplant recipients receiving a A*01:01 mismatched allograft. Data shown as the percent of recipients (y axis) with a 2MM residue located at positions 1 to 274 of the A*01:01 α-chain (x axis). 2MM residues within the AH-PBG region are highlighted in the teal boxes. **(K)** Histogram showing frequency distribution and location of solvent-accessible (250 Å^2^) 2MM residues within the A*01:01 epitopes recognized by E07 and L02 rmAbs in 11 transplant recipients who received a mismatched A*01:01 allograft. The total number of solvent-accessible 2MM residues located within the AH-PBG region was determined for each subject and the percentage of these 2MM residues that were contained within the A*01:01 epitopes bound by E07 and L02 was determined. **(L-M)** Chimeric mouse/human rmAbs cE07 and cL02, derived from Bmem isolated from the kidney of recipient N006, block binding to A*01:01 by polyclonal Abs found in serum samples isolated from 11 DSA^+^ transplant recipients who received a A*01:01 mismatched allograft. Data reported as percent inhibition of serum binding and shown as the mean two independent experiments with at least two technical replicates/experiment. W6/32, a murine Ab specific for human class I HLA, included as a negative control for inhibition. **(N)** Classification of HLA α-chain residues 158, 163, and 166 for 11 DSA^+^ transplant recipients who received a mismatched A*01:01 allograft. The residues present in A*01:01 at positions 158, 163 and 166 were compared to the residues expressed by each subject’s self-HLA-A alleles at the same position. Residues were classified as 2MM, 1MM or 0MM between A*01:01 and the self-HLA alleles. **(O-P)** Binding to WT A*01:01 and mutant A*01:01 (V158A, R163T, D166E) by serum samples from 11 DSA^+^ transplant recipients who received a A*01:01 mismatched allograft. Data shown as binding activity (arbitrary units, AU) for each WT or mutant A*01:01 protein (**O**) or as percentage of WT binding activity in **(P)**. Binding activity was calculated by interpolating binding activity using a standard curve of W6/32 binding for each mutant (see *Methods* for details). Data are shown as mean values from three independent experiments for each subject:protein combination. **(Q)** Correlation between loss of binding to HLA-A*01:01 mutants (V158A, R163T, D166E) and chimeric cE07-dependent binding inhibition in serum samples from 11 DSA^+^ transplant recipients who received a A*01:01 mismatched allograft. Data shown as scatterplots with x-axis displaying % inhibition by chimeric cE07 (top plot) or cL02 (bottom plot) rmAbs and y-axis displaying percent binding (mutant binding AU/WT binding AU × 100). Values in (**A,B**) were compared against the non-blocking condition using the Kruskal-Wallis test with Dunn’s multiple comparisons test. Values in (**D**) were compared against WT binding using the Kruskal-Wallis test with Dunn’s multiple comparisons test. Proportions in (**G**) were compared using Fisher’s exact test. Group ranks in (**L**) were compared using the Friedman test with Dunn’s multiple comparisons test against the W6/32 condition. Group ranks in (**O**) were compared using the Friedman test with Dunn’s multiple comparisons test against the WT protein. Statistical analysis in (**Q**) performed using a nonparametric Spearman correlation with a two-tailed P value. See Supporting Figures and Tables for additional A*01:01 epitope data for the n=247 US cohort and the n=11 SOT cohort (Figure S6, Table S1).

### Shared recognition of immunodominant HLA-A epitopes across transplant recipients

Our data showed that recipient N006 made a serologic response to the A*01:01 mismatched allograft that could be almost completely inhibited with two rmAbs, which targeted the A*01:01 AH-PBG region and were derived from the N006’s kidney Bmem (Figure 7F). While the AH-PBG region is comprised of 74 residues, or 27% of the entire 274 AA A*01:01 α-chain (Figure 7G), we found that 73% of the 41 epitopic residues recognized by E07 and L02 were localized within the AH-PBG region (Figure 7G). As this frequency was significantly greater (p<0.0001) than would be expected by chance, the data suggested that the topography of the HLA-A protein was critical for driving this focused B cell response. To address this idea, we next asked whether new B cell epitope prediction models, such as GraphBepi^58^ and DiscoTope-3.0^59^ that incorporate machine learning via the quaternary protein structure prediction tool AlphaFold2, could more accurately identify these immunodominant topographically defined allo-HLA epitopes. Interestingly, both tools predicted that most of the B cell response against A*01:01 would focus on the exposed AH-PBG residues (Figure 7F). In fact, 86% and 80% of the B cell epitopic residues predicted by GraphBepi and DiscoTope-3.0, respectively, were contained within the AH-PBG region (Figure 7G) – a frequency that was again significantly higher (p<0.0001) than expected by chance.

The computational prediction tools supported our experimental data showing that the B cell and serologic response of recipient N006 was focused on the AH-PBG region of A*01:01. Since the combination of the E07 and L02 rmAbs covered 41% (30/74) of the residues and 54% of solvent-accessible surface area in the A*01:01 AH-PBG region for recipient N006, we hypothesized that for any given individual, more than half of any solvent-accessible 2MM residues present in the AH-PBG would be encompassed within the A*01:01 epitopes recognized by E07 and L02. To assess this possibility, we mapped the location of all residues in a cohort of 247 HLA-A genotyped subjects who did not express A*01:01 as one of their two self-HLA-A alleles. We determined how many of the residues were solvent-accessible, double-mismatched between non-self (A*01:01) and the participant’s self-HLA-A alleles, localized within the AH-PBG region and contained within the A*01:01 epitopes recognized by E07 and L02 rmAbs. For most subjects, we identified a median of five solvent-accessible 2MM AA residues that were localized within the AH-PBG region (Figure S6E). Interestingly, most of these 2MM residues were also encompassed within the epitopes recognized by E07 or L02 (Figure 7H). Indeed, for 73% (180/247) of the subjects in this representative cohort, 100% of the A*01:01 AH-PBG region localized, solvent-accessible 2MM residues were contained within the A*01:01 epitopes bound by the E07 and L02 rmAbs (Figure 7H).

Our computational data supported the concept that allo-A*01:01 specific B cell and serologic Ab responses could converge on A*01:01 epitopes that are localized within the AH-PBG region and potentially “covered” by the E07 and L02 rmAbs. If correct, we hypothesized that chimeric cE07 and cL02 rmAbs should block binding of DSA collected from other transplant patients who generated a serologic response to a mismatched A*01:01 allograft. We further postulated that the inhibition mediated by cE07 and cL02 rmAbs should not be dependent on the individual’s self-HLA. To experimentally test this hypothesis, we examined serum collected from 11 transplant recipients who received a mismatched A*01:01-typed organ and subsequently developed circulating anti-A*01:01 DSA. As expected, all 11 individuals made a robust anti-A*01:01 response to the mismatched alloantigen (Figure 7I). However, they also developed unique reactivity patterns to third-party HLA (Figure 7I). Consistent with our earlier analysis of recipient N006 (Figure 5L), about 62% of the 98 solvent-accessible 2MM residues in A*01:01 that were identified across all 11 transplant recipients were located within the A*01:01 AH-PBG region, with the remaining 38% of the 2MM residues located outside of the AH-PBG region (Figure 7J). In addition, we noted that the epitopes recognized by E07 and L02 encompassed every 2MM residue within the AH-PBG for 63% (7/11) of the subjects (Figure 7K, Figure S6F) – a number that was similar to the percentage calculated for the representative national cohort (Figure 7H). Next, we used cE07 and cL02 rmAbs to test whether the circulating allospecific Ab present in these 11 A*01:01-mismatched recipients was focused on the AH-PBG region of A*01:01. Strikingly, we observed that cE07 and cL02 rmAbs alone and in combination significantly inhibited (p<0.0001) A*01:01 binding by circulating alloAbs from the serum of all 11 transplant recipients (Figure 7L). Indeed, the combination of chimeric E07 and L02 rmAbs inhibited 83-100% (mean 97%) of A*01:01-binding by circulating serum alloAbs present in the 11 transplant recipients (Figure 7M). Importantly, the W6/32 Ab, which does not bind to the AH-PBG region, showed only 5.2% average inhibition of binding by the serum from these same recipients (Figure 7M).

To address whether binding of the A*01:01-specific polyclonal Abs present in these 11 transplant patient serum samples was dependent on 1MM or 2MM residues localized within the AH-PBG region we compared binding of the subjects’ polyclonal serum alloAbs to WT A*01:01 and the 3 A*01:01 mutants V158A, R163T and D166E. Notably, even though the 11 transplant recipients expressed different combinations of self-HLA-A molecules (Figure 7I), 11/11 of the transplant recipients were double-mismatched at V158, 9/11 of the recipients were double-mismatched at R163, and 6/11 of the recipients were double-mismatched at D166 (Figure 7N). Consistent with our hypothesis that the B cell and serologic response would be focused on the mismatched residues localized within the AH-PBG region, the serum Abs from these recipients bound less well to the 3 mutant A*01:01 proteins relative to WT A*01:01 (Figure 7O). Indeed, while not significantly different across the entire cohort (Figure 7O), binding of the serum polyclonal Abs to V158A was reduced by an average of 32% (Figure 7P). More strikingly, polyclonal alloAb binding to R163T and D166E was significantly decreased (Figure 7O) with an average reduction in binding of 57-61% across the 11 individuals (Figure 7P). When we then compared inhibition by cE07 with changes in binding to the A*01:01 mutants, we observed that serum samples that were highly inhibited by cE07 bound significantly less well to the R163T and D166E A*01:01 mutants (Figure 7Q). By contrast, there was no correlation between inhibition by cL02 and sensitivity to the A*01:01 mutants that targeted mismatched residues present in the epitope recognized by E07 (Figure 7Q). Thus, the cE07 rmAb appeared to inhibit binding of polyclonal alloAbs from the 11 transplant recipients by binding to shared, specific epitopic residues located within the AH-PBG region. Taken altogether, these data suggest that the serologic response to mismatched A*01:01 made by 12 different transplant recipients is highly focused on immunodominant epitopes that are contained within the solvent-exposed residues that are localized within the AH-PBG topographic region. We further showed that two rmAbs, which were cloned from kidney-residing B cells in one recipient, not only targeted these immunodominant HLA-A epitopes in that individual but also potently inhibited binding of A*01:01 DSAs from completely unrelated transplant recipients. Therefore, we argue that the tissue and circulating allospecific Bmem compartment is focused on the same immunodominant “public” HLA epitopes and is tightly linked to the pathogenic serologic response toward the HLA alloantigens. The relevance of these findings to HLA matching strategies and to potential immunotherapies directed against humoral alloresponses is discussed.

## DISCUSSION

Despite the fact that anti-HLA DSA^2,4^ and HLA-specific B cells^7–9^ are clearly associated with organ transplant rejection,^9^ the rules governing the recognition of HLA alloantigens by human B cells are still largely unknown. While extensive analyses of the HLA alloreactivity patterns generated by polyclonal DSA present in transplant recipient sera have been performed, these patterns cannot be easily deconvoluted to understand the specific reactivity profile of individual allospecific B cells or individual alloreactive Ab molecules.^60^ Therefore, to understand the fine specificity of the anti-HLA alloreactive human B cell response and the immunogenicity of nonself HLA, the HLA-alloreactive human B cell response needs to be studied at the single cell level with a focus on the structural basis for B cell allorecognition. However, to date, only a single structure containing HLA and an anti-HLA Ab, the latter of which was generated via phage-display, has been solved.^31^ Likewise, single cell BCR repertoire and specificity analyses of HLA-specific human B cells have been limited to small numbers of rmAbs.^26,61^ Here, we cloned and expressed 50 BCRs derived from a transplant recipient’s kidney and blood B cells that were specific for the mismatched A*01:01 expressed by the donor allograft. Using these allospecific rmAbs, we showed that the anti-HLA B cell response was affinity-matured, positively selected, shared across kidney and blood, and clonally linked to the clinically important circulating DSA. We identified three clonally distinct lineages of allospecific B cells that collectively contributed to over 70% of the alloreactive rmAb repertoire and used representative rmAbs from each of these immunodominant lineages to assess the structural basis for B cell allorecognition. While AA residues that are mismatched between donor and self-HLA can be found throughout the entire 274 AA HLA-A protein, we found that the B cell response of this individual was focused on a small subset (∼27%) of the solvent-accessible, topographically exposed, mismatched residues present in the A*01:01 α1 and α2 helices that form the crown of the HLA molecule and frame the peptide binding groove (AH-PBG region). This focusing of the B cell response on the HLA crown or AH-PBG region did not appear due to chance, suggesting that this topographically defined region was selectively targeted by the recipient’s B cells and Abs. Surprisingly, this immune focusing on the HLA crown was not limited to a single individual as an additional validation cohort of transplant recipients who also received an A*01:01-mismatched allograft generated immunodominant anti-donor responses that targeted the same topographic region of HLA. Moreover, even though these individuals expressed a diverse array of self-HLA alleles, the circulating DSA from these subjects was uniformly sensitive to mutations in the AH-PBG region, indicating the convergence of these alloresponses upon the topographically exposed crown of the HLA molecule. These data therefore argue that specific mismatched residues are part of “public” B cell allo-epitopes and are targets for immunodominant, convergent B cell and Ab responses across a range of individuals.

Our data suggests that the combination of mismatched residues within the quaternary structure of the allo-HLA-A antigen dictates B cell recognition of the non-self allele. This is different from one of the leading current theories in the field, referred to as eplet recognition.^26,30,60,62,63^ In the eplet model, mismatched residues are grouped into small, 3.5 Å-radius patches of mismatched residues called eplets that are hypothesized to be recognized by the heavy chain CDR3 (HCDR3) of alloreactive Abs. Our structural and reactivity data for the 50 rmAbs cloned from the blood and kidney of a transplant recipient undergoing AMR did not easily fit the eplet model. First, we found that almost half (44%) of these rmAbs could not be matched to any known eplet-based putative pattern of reactivity. Second, our structural analyses of A*01:01 complexed with E07, M07 and L02 revealed disconnects between predicted eplets and the actual documented physical interactions between these alloAbs and HLA. Third, both the IgH and IgL chains from all three rmAbs provided critical contacts with the HLA residues that contribute to HLA binding. Finally, we found that the key specificity and reactivity-conferring HLA residues within the epitope recognized by the allospecific rmAbs could be considerably farther apart than 3.5 Å. These findings, while in opposition to the eplet model, were entirely consistent with our broad understanding of B cell epitopes, as studies of hundreds of Ab/antigen structures have shown that most epitopes are approximately 1100 Å^2^ and that the HCDR3 is not the only Ab subdomain that is important for the fine specificity of a given Ab.^15,48,64^ Thus, we argue that HLA epitope prediction programs may potentially benefit from a more focused examination of the 1MM and 2MM residues that are solvent accessible and located within the AH-PBG region and that distance of mismatched residues to the center of the PBG might a useful metric to identify the most immunogenic mismatched residues in HLA-A.

Our data suggests that specific regions of the alloHLA-A molecule are selectively and repetitively targeted by the B cell and humoral immune response. Indeed, we documented 8 distinct A*01:01-monospecific clonal/genetic lineages that targeted the same or overlapping epitope in recipient N006. Although multiple factors contribute to B cell immunodominance,^17^ recent work has increasingly emphasized the importance of antigen orientation to the fine specificity of antiviral responses.^52,65–67^ In order for BCRs to bind cognate antigen, that antigen must first be physically accessible; thus, antigenic topography acts as an initial checkpoint for epitope selection.^65^ Given that the topographically exposed crown of the HLA molecule serves as the binding site for other immune receptors, including the T cell^68^ and some NK cell receptors,^69^ we argue that the exposed crown of HLA also supports recognition and binding by BCRs that target solvent-accessible mismatched residues. Convergent epitope targeting is frequently observed in other settings – including responses to SARS-CoV-2,^70,71^ influenza,^72^ and Ebola.^73^ In our study, we demonstrated epitopic convergence within both the B cell and serologic arms of the humoral response in a single individual, as well as epitopic convergence across a cohort of transplant recipients. Indeed, in this larger cohort of transplant recipients, a cocktail of two rmAbs that only buried 11.4% of the entire A*01:01 α-chain surface (E07 and L02 epitopes = 1,493 Å^2^; entire A*01:01 α-chain surface area = 13,117 Å^2^) was nonetheless able to inhibit on average 95% of circulating DSA binding by targeting common, “public” immunodominant epitopes within the AH-PBG region. Importantly, this immune focusing occurred regardless of the transplant recipients’ self-alleles, suggesting that immune focusing is a general feature of the response to nonself HLA. In the context of antiviral responses, it is appreciated that AA polymorphisms contribute to immune focusing and immunodominance.^19^ Although various methods have been used to explain why certain HLA residue mismatches may be immunogenic, including hydrophilicity,^74^ protrusion,^51^ and other physicochemical features,^49,50,75^ these approaches have all generally lacked discriminatory power. Indeed, none of these physicochemical features of polymorphic mismatched residues correlated with immunogenicity in our study. Instead, our data argued that the topography of the antigen itself and the accessibility of mismatched residues within a discrete region of the quaternary structure dictate immunogenicity. In transplantation, this observation of topographic immunodominance has several implications. First, it provides a framework for prioritizing HLA mismatching based upon the topography of mismatched residues. Second, it suggests that targeted therapeutic approaches that either block or mutate these highly immunogenic crown residues might prove effective in abrogating alloantibody responses in transplant recipients. In addition, whereas most prior studies of antigen topography have relied on mouse models^52,66^ or strictly computational models,^65^ our data shows the contribution of topography to ongoing human B cell immune responses.

Human allotransplantation presents an immune problem with several unique features relative to other immune contexts – namely the presence of a foreign antigen that is highly similar to self and the cover of ongoing pharmacologic immunosuppression. Nonetheless, we observed many expected features of a canonical, “high quality” adaptive immune response – suggesting the application of these alloresponse findings to broader contexts. Clearly, the chronic nature of transplant rejection and the antigenic volume should support robust clonal selection and affinity maturation – consistent with observations in infection,^76^ autoimmunity,^40,77^ and cancer.^78^ However, although other studies^79–81^ in transplantation have demonstrated clonal expansions in the B cell repertoire; there is only relatively limited data^27^ showing that these clonally expanded populations specifically target HLA. Our study shows that the anti-donor HLA B cell response in both the kidney and blood can be affinity matured and clonally selected for recognition of specific immunodominant epitopes. These clonally selected lineages were composed of multiple B cell and ASC subsets that spanned the blood and kidney. Indeed, the three rmAbs that were used for the structural studies and were representative of the three immunodominant epitope recognition groups were each cloned from kidney-infiltrating Bmem cells. Tissue-infiltrating B cells have been demonstrated to show antigen specificity in a variety of human contexts – including transplantation,^79^ cancer,^78,82^ and infection.^83^ Like other Bmem cells, tissue-residing Bmem cells have the potential to rapidly differentiate into ASCs that secrete Ab locally within the tissue. Our data showing many clonal connections between the allograft-infiltrating B cells, the B cells in circulation, and the serologic pathogenic DSA suggest that the tissue-residing Bmem have the potential to replenish the ASC pool that can secrete pathogenic Abs directly within the allograft and contribute to long-term systemic DSA responses after migrating to the bone marrow. Bmem cells within lymphoid and non-lymphoid tissues can also contribute to Ab-independent effector functions like cytokine production and antigen presentation. Given the highly diverse phenotype of intragraft B cell compartment, which includes Bmem and DN B cells, we expect that these graft-infiltrating cells may influence the immune microenvironment within the graft. Although we were unable to identify tertiary lymphoid follicles within the allograft, we did find clusters of proliferating B cells that were in close proximity to T cells, raising the possibility that the allospecific B cells might interact with intragraft T cells. However, our data does not resolve whether the allospecific B cells found in the allograft were generated *in situ* or migrated to the tissue following affinity maturation in secondary lymphoid tissues. Regardless, we argue that the allograft can serve as a reservoir for multiple diverse clones of alloreactive B cells and ASCs.

In conclusion, this work provides a framework for understanding the structural and molecular basis of B cell epitope targeting in organ transplant rejection. It highlights the anatomy of this B cell alloresponse as it spans tissue and blood and results in the generation of a high-affinity donor HLA-specific response that can be detected as circulating DSA. Our data indicate that topography and quaternary structure of HLA-A enforces immune focusing and immunodominance for B cell responses in transplant rejection. More broadly, these data support the surprising conclusion that B cell immunodominance hierarchies in alloimmune responses are driven by the orientation and topography of the HLA antigen and that this process of immunodominant immune recognition is not entirely dependent on the constellation of self-HLA-A proteins expressed by the recipient. These findings therefore may in the future allow for a more precise identification of immunogenic “public” residues in non-self HLA, which in turn could lead to better algorithms for organ matching. In addition, the findings suggest that it may be possible to intervene or prevent AMR using a collection of Ab-based biologic blocking agents that are specifically designed to recognize and cover the immunodominant epitopes in the AH-PBG region on the crown of the HLA protein. Finally, the molecular and structural data associated with B cell alloresponses in transplant recipients provide us with the unique opportunity to establish a more nuanced understanding of the fundamental processes of tolerance and immunity in humans.

### Limitations of the study

One limitation to this study is that we only studied Ab responses to the mismatched HLA and it is certainly possible that Abs specific for other non-HLA antigens expressed by the transplanted organ may contribute to transplant rejection.^84^ However, it is well appreciated that HLA-specific Abs are very important players in organ transplant rejection^9^ and that understanding the rules governing non-self HLA recognition by B cells has the potential to improve transplant outcomes. A second limitation is that most molecular, proteomic, and structural experiments were conducted with samples from a single individual (N006) who underwent a kidney nephrectomy following rejection of a second transplanted kidney. Despite this, the depth of sampling of single allo-HLA specific B cells was substantially greater than any previous study,^26^ and the thousands of A*01:01-reactive IgV_H_ sequences, 14 distinct A*01:01-reactive lineages, and three dominant specificities indicate that we examined a robust yet diverse alloimmune response in this individual. Moreover, our proteomic and molecular data allowed us to, for the first time, directly link the intragraft alloimmune response to the serologic pathogenic DSA response. Most importantly, the serologic features of immunodominance observed in recipient N006 appeared to be a general phenomenon that was maintained within a larger cohort of transplant recipients expressing distinct self-HLA-A alleles who also appeared to mount a shared convergent response to the donor-specific A*01:01 protein expressed by their mismatched allografts. A final limitation of this study is that we did not assess the contribution of T cell help to HLA epitope focusing by the B cells.^17^ However, we did confirm that the allo-HLA rmAbs bound to A*01:01 expressed on plasma membrane of cells, suggesting that Ab recognition of the crown of the HLA-A protein is not dependent on loading of a specific peptide. Going forward, it will be important to employ similar single cell approaches to examine B cell alloHLA immune responses in other individuals who exhibit rejection pathologies involving other mismatched HLA proteins, like HLA class II, which is commonly targeted by the alloreactive B cells in late transplant rejection.^4^ While it is possible that there will be differences in the T and B cell responses to mismatched HLA based upon the class or locus of the mismatched HLA, we speculate that the HLA immunogenicity “rules” will be similar given the structural and topographic similarities between class I and class II molecules.^85^

## ACKNOWLEDGMENTS

We acknowledge the NIH Tetramer Core facility for providing HLA-A*01:01, HLA-A*02:02, and HLA-A*24:02 tetramers, monomers, and expression plasmids and the Southeast Regional Collaborative Access Team (SER-CAT) for access to the 22-ID-D beamline at the Advanced Photon Source, Argonne National Laboratory. We thank Vidyasagar Hanumanthu, director of the UAB Flow Cytometry and Single Cell core, and Dr. Michael Crowley, director of the UAB Heflin Genomics core, for their assistance with these studies. We thank the CryoEM facility of La Jolla Institute for Immunology, as well as philanthropic support of this facility.

## AUTHOR CONTRIBUTIONS

F.E.L. and J.T.K. conceived the study. F.E.L. secured funding, oversaw project administration, participated in the original drafting of the manuscript and its revisions, and provided direct supervision of the overall project. J.T.K. conducted experiments, performed formal analyses, curated data, visualized data, wrote the original draft of the manuscript and participated in its revisions. R.G.K. conducted experiments, provided methodologic expertise, validated key reagents, conducted formal analyses, curated data, assisted with proteomic visualization, provided supervision and participated in the drafting of the manuscript and its revisions. A.C.K.L. conducted experiments, performed formal analyses, and participated in revision of the manuscript. T.J.G. provided methodologic expertise, conducted experiments and formal analyses, curated data, participated in the drafting of the manuscript and its revisions, and provided supervision. J.L.K conducted formal analyses of structural data. C.F.F. provided methodologic expertise, conducted formal analyses, and generated visualizations of repertoire data. A.F.R provided methodologic expertise and supervision. R.D.A, S.Q., A.S.S., and B.J.M. conducted experiments. K.J.M., A.R.C., G.Y., M.E.H., and J.A. provided key reagents. D.B.G., S.K., S.C.O., V.K., J.A.H., and F.D.R assisted with formal analyses. J.F.K. and T.D.R supervised staff associated with this project. E.O.S supervised, facilitated project administration, and provided critical resources. P.M.P. reviewed and edited the manuscript. All authors reviewed and approved the final version of the manuscript.

## METHODS

### Subjects

A kidney transplant recipient (subject N006) with a history of allograft rejection and graft loss who was undergoing transplant nephrectomy was enrolled in this study at the University of Alabama at Birmingham (UAB) (Table S1 for clinical history). This study was approved by the UAB Institutional Review Board (IRB, protocol # IRB-300004168), and informed consent was provided by the study subject. Serum samples from n=11 solid organ transplant recipients (Table S1) who received an HLA A*01:01-mismatched solid organ transplant and subsequently developed anti-A*01:10 DSA were analyzed. This study was approved by the Institutional Review Board at UAB (protocol # IRB-300007045), and informed consent was provided by the study subjects. Biologic samples from a single deceased organ donor, deemed non-human subjects research by the UAB IRB, were used.

### Clinical anti-HLA Ab testing

Clinical anti-HLA Ab testing for recipient N006 and the cohort of n=11 solid organ transplant recipients was performed by the UAB Histocompatibility Laboratory using its standard clinical assay. For the *clinical* MFI data shown in Figure 1I, Figure 7I, and Table S1, IgG Abs from patient sera were collected using a Melon IgG spin column and then analyzed neat using the OneLambda HLA arrays, according to manufacturer instructions (OneLambda, ThermoFisher).^87^ Proprietary software was used for the MFI calculation with a MFI ≥5,000 used as the clinical threshold for a positive reading. HLA-specific Ab data was obtained from the patients’ electronic medical record. All *research* HLA-Ab cytometric bead array data were generated in the research lab using methods described below.

### Clinical HLA typing by next-generation sequencing or by imputation

HLA typing of both donor (donor to recipient N006) and recipient (subject N006) was performed by the UAB Histocompatibility Laboratory. Genomic DNA was extracted from peripheral blood samples using the Qiagen EZ1 DNA blood kit via the Buffy coat protocol. Library construction and enrichment were performed per manufacturer’s protocols for the CareDx AlloSeq Tx 17 assay and libraries were run on the Illumina MiSeq. Propietary CareDx software, AlloSeq Assign, v1.0.3 (CareDX, Stockholm, Sweden) utilized references from the Immuno Polymorphism Database-ImMunoGeneTics project/HLA Database (IPD-IMGT/HLA database) v3.47.0.0 to make allele assignments for all HLA Class I and Class II loci. For donors and recipients in the n=11 cohort that were not HLA typed by next-generation sequencing, 4 digit HLA-A alleles for recipient and/or donor were imputed (Table S1) based upon ethnicity and population frequencies.^88^

### Immunofluorescent histology of allograft specimens

Tissue sections (6 µm) from the transplant nephrectomy specimen were embedded in Tissue-Tek O.C.T. Compound (Sakura), frozen in LN_2_ and stained with three anti-human Ab panels including CD21::FITC (Biolegend, clone Bu32), CD19::APC (BD, clone HIB19), CD21::FITC (Biolegend, clone Bu32), CD4::AF488 (Biolegend, clone OKT4), CD19::PE (Biolegend, clone HIB19), Ki67::APC (Biolegend, clone Ki-67), anti-Ig::AF488 (Southern Biotech, polyclonal), CD38::APC (Biolegend, clone HIT2), and DAPI. Images were acquired on a Nikon Eclipse Ti microscope with the eyepiece at 10x, mirror at 1x, and objective at either 20x or 40x (total 200x or 400x magnification). Original images including metadata and exposure times saved as *.ND2 files then exported at 300dpi as *.TIFF files. Exposure times (in milliseconds) for each channel (DAPI, FITC, PE, APC): Figure 1A (500, 3000, 6000, 10000), Figure 1B (2000, 3000, 8000, 10000), Figure 1C left (60, 70, 1000, 600), Figure 1C right (250, 70, 1400, 800). The tonal range of RGB was reduced in Adobe Photoshop (Adobe Inc., San Jose, CA) to increase brightness and contrast, with settings uniformly applied across all images utilizing the same staining panel.

### Multi-color flow cytometry and sorting

The allograft nephrectomy specimen was mechanically and enzymatically digested to yield a single cell suspension. Mononuclear cells were isolated from peripheral blood using density-gradient centrifugation. The cell suspensions were stained with the following anti-human Ab reagents: CD19::V500 (BD, clone HIB19), CD71::BV605 (BD, clone M-A712), CD38::BV786 (BD, clone HIT2), IgD::FITC (BD), CD11c::PE (Biolegend, clone Bu15), CD27::PE-CF594 (BD, clone M-T271), 7-AAD (BD), CD3::PE-Cy5 (Biolegend, clone HIT3a), CD14::PE-Cy5 (Invitrogen, clone 61D3), CD16::PE-Cy5 (BD, clone 3G8), CD56::PE-Cy5 (BD, clone B159). Cells were also stained with fluorescent-labeled (BV421 and AF647) tetramers of recombinant HLA-A*01:01 (NIH tetramer core, loaded with peptide VTEHDTLLY). Cells were sorted on a FACSAria (BD Biosciences) in the UAB Comprehensive Flow Cytometry and Single Cell Core as single cells for recombinant monoclonal expression and in bulk populations for next generation IgV_H_ sequencing. Data were analyzed using FlowJo v10.8.1 (BD).

### Next generation sequencing of the IgV_H_ repertoire (IgV_H_-Seq)

IgV_H_ sequencing was performed as previously described.^86^ After sorting B cells subsets directly into lysis buffer (Norgen Single Cell RNA Purification Kit) with 1% mercaptoethanol, lysates were snap frozen on dry ice and stored at −80°C. Total cellular RNA was extracted from bulk sorted B cell and ASC subsets according to the manufacturer’s protocol. cDNA was prepared (BioRad iScript) per manufacturer’s instructions, and DNA amplicon products were generated using Ig-specific variable and constant gene primers.^86^ Samples were indexed (Nextera Index Kit, Illumina), purified, quantitated (KAPPA Library Quantitation Kit, Roche) and pooled into libraries. Following generation, libraries were denatured per manufacturer instructions (Illumina) and loaded onto a 600-cycle V3 MiSEQ cartridge (Illumina) for sequencing^40^ in the UAB Heflin Sequencing Core.

### Phylogenetic analyses of IgV_H_-Seq data

IgV_H_ sequencing data were processed and analyzed as described previously.^40^ Joined paired-end reads were assembled and quality filtered using FastQC scores. Using IMGT/HighV-QUEST,^89^ sequences were annotated, and productive sequences were utilized for downstream analyses. Sequences were clustered into lineages based upon *IGHV* and *IGHJ* gene identity, identical HCDR3 length, and HCDR3 nucleotide identity >85%. Downstream analyses were performed in Matlab (R2020a, The Mathworks Inc.) or in R (R core team). Phylogenetic trees were constructed using Phylip’s DNA parsimony (dnapars) tool (v3.695; adjusting settings 1, 4, 5, 6 and O setting the germline sequence as the outgroup).^90^ Phylogenetic trees were visualized using Cytoscape v3.8.2.^91^

### Circos visualization of IgV_H_-Seq data

The outer numbering of Circos plots shows sequence counts for a given tissue-phenotype cell subset. The outer ring colored (non-gray) for lineages contained in the top 20% of size-ranked lineages within a given tissue-phenotype cell subset. Links connect lineages shared between tissue-phenotype subsets; links are colored (non-gray) if the lineage is contained in the top 20% of size-ranked lineages in at least one tissue-phenotype cell subset. Alluvial ribbons depict connectivity of shared lineages, with the individual lineages ranked by size in each subset along the y-axis. Shared lineages are represented as ribbons with individual lineages ranked by size in each subset. The numbers of sorted cells for IgV_H_ sequencing, numbers of unique non-singleton lineages and %D50 are indicated. %D50 = (# lineages in top 50% of subset A / # of lineages in the top 50% of subset B) × 100.

### Single cell rmAb cloning, sequencing, expression and purification

rmAbs were generated using previously described methods.^86^ cDNA was prepared from mRNA isolated from single non-naïve (CD19^+^IgD^neg^) dual HLA-A*01:01 tetramer-binding B cells index-sorted into hypotonic lysis buffer in 384-well PCR plates which were stored at −80°C until used. RT-PCR was performed to amplify IgV_H_ and IgV_K/L_ transcripts. Amplicons were cloned into human IgG1, IgK, or IgL expression vectors, as previously described.^86^ The resulting plasmids underwent Sanger sequencing at the Genetic Resources Core Facility (RRID SCR_018669, Johns Hopkins University Department of Genetic Medicine, Baltimore, MD). IgH and IgL plasmids for each Ab were co-transfected into 293FreeStyle cells (Invitrogen) using polyethylenimine (Polysciences). Conditioned supernatants were screened for anti-HLA reactivity using HLA cytometric bead arrays as described below. HLA-A*01:01 reactive rmAbs were purified from conditioned media using Sepharose G Fast Flow (Cytiva) and the resulting rmAb concentrations were determined by UV spectrophotometry (Nanodrop, Thermo Fisher). For x-ray crystallography and cryoEM studies, the heavy chain expression vector was modified using site-directed mutagenesis to remove the IgC_H_ gene segment of each rmAb (E07, L02, and M07). The resulting constructs were transfected into 293Freestyle cells as described above to produce antigen-binding fragments (Fabs). Fabs were purified using CaptureSelect IgG-CH1 Affinity Matrix (Thermo Scientific).

### Generation of modified and UCA rmAbs

UCAs for each rmAb were generated by reverting all nucleotides encoding variable gene segments (V_H_, D_H_, J_H_, V_K/L_, J_K/L_) to the IMGT reference sequence of the gene segment alleles. Nontemplated nucleotides (N-additions) were not modified. Recombinant human IgG1 mAb constructs were synthesized (Sino Biological) using the predicted UCA nucleotide sequences.

### Immunogenetic analysis of rmAbs

IgV_H_ and IgV_K/L_ sequences from the cloned and verified A*01:01-specific rmAbs were submitted to IMGT/HighV-Quest for annotation. rmAbs were grouped into clusters based on identical V_H_, D_H_, J_H_, V_K/L_, J_K/L_ gene assignments and identical HCDR3 and LCDR3 lengths. All 50 rmAb IgV_H_ sequences were then directly compared to the sequences in the bulk-sorted kidney and blood B cell/ASC IgV_H_-Seq database. Any rmAb meeting the lineage criteria (same VH, DH, JH gene assignments, identical HCDR3 length and >85% NT identity across the HCDR) were assigned as members of the bulk B/ASC clonal lineages and these lineages were defined as A*01:01 specific. This approach resulted in the identification of 14 unique A*01:01-specific lineages, which each contained one or more A*01:01-specific rmAbs (see Table S2).

### High-throughput localized surface plasmon resonance (L-SPR)

L-SPR data were collected using a 1:1 referencing system on the Nicoya Alto HT-SPR instrument according to manufacturer’s recommendations and as described.^92^ Biotinylated recombinant WT HLA-A*01:01 (500 µg/mL, NIH Tetramer core or produced in-house) or single residue mutants of HLA-A*01:01 (500 µg/mL, produced in-house) was resuspended in PBS containing 0.05% Tween-20 and immobilized on an EDC/NHS-activated carboxyl sensor (Nicoya) for 20 min. The rmAbs (analytes) were diluted to 1.3 µM in PBS containing 0.05% Tween-20. Single-cycle kinetic analyses were performed at 25°C with five threefold analyte dilutions, with maximum concentrations of 433 nM. Kinetic fitting was performed using a 1:1 Langmuir model within the Nicoya user portal. For datasets that poorly fit with a 1:1 Langmuir model, an affinity fit model was used (TraceDrawer, Ridgeview Instruments). The limit of detection for the equilibrium constant (K_D_) was estimated to be 4.3 µM (10 times the highest concentration of analyte tested). Recombinant mAbs with responses <50 plasmon resonance response units or without monotonic increases across increasing concentrations were judged to not have measurable binding by L-SPR. For visualization and for calculations of fold-change in affinity, if a rmAb had no detectable binding by L-SPR (K_D_ >4.3 µM), a K_D_ of 5 µM was used.

### Plasma immunoglobulin proteomics

Recombinant HLA-A*01:01 was conjugated to CNBr-activated Sepharose (Cytiva) following manufacturer’s protocol. Recipient N006 plasma was diluted 1/10 in PBS, filtered using a 22mm syringe filter (Sigma-Aldrich) and passed over the HLA-A*01:01 column. Bound protein was eluted with 0.1M glycine pH 2.5, buffer-exchanged into PBS, concentrated to 1mg/mL, and stored at 4°C. The resulting affinity-purified A*01:01-binding polyclonal plasma Abs (appAbs) were reduced with DTT, denatured at 70°C for 10 minutes (min) and separated on a 10% Bis-Tris Protein gel. IgH and IgL bands were visualized with colloidal Coomassie, excised from the gel, equilibrated in 100 mM ammonium bicarbonate (AmBc), and digested overnight with Trypsin Gold, Mass Spectrometry Grade (Promega) or endoproteinase AspN (New England BioLabs) following manufacturer’s instructions. Peptide digests were reconstituted in 0.1% Formic Acid/ddH_2_O at 0.1 µg/µL, injected (8 µL each) onto a 1260 Infinity nHPLC stack (Agilent Technologies), and separated using a 75 µm I.D. x 15 cm pulled tip C-18 column (Jupiter C-18 300 Å, 5 µm, Phenomenex). This system runs in-line with a Thermo Q Exactive HFx mass spectrometer, equipped with a Nanospray FlexTM ion source (Thermo Fisher Scientific). Data were collected by the UAB Mass Spectrometry / Proteomics Core in CID mode. The nHPLC was configured with binary mobile phases that includes solvent A (0.1% FA in ddH2O), and solvent B (0.1% FA in 15% ddH2O / 85% ACN), programmed as follows: 10 min at 5% solvent B (2 µL/min, load), 90 min at 5%-40% solvent B (linear: 0.5 nL/min, analyze), 5 min at 70% solvent B (2 µL/min, wash), 10 min at 0% solvent B (2 µL/ min, equilibrate). Following each parent ion scan (300-1200m/z at 60k resolution), fragmentation data (MS2) was collected on the top-most intense 10 ions at 7.5k resolution. For data dependent scans, charge state screening and dynamic exclusion was enabled with a repeat count of 2, repeat duration of 30 seconds (s), and exclusion duration of 90 s. Following LC-MS/MS, the data were processed, searched, filtered, grouped, and quantified, as previously reported.^93^ LC-MS/MS-derived peptide AA sequences were used to probe an AA sequence database that included all IgV_H_ sequences derived from the kidney, blood and rmAb IgV_H_-seq libraries. Exact peptide matches to IgV_H_ sequences were determined and members of the lineages identified were ranked by % of total AA matched. For visualization, peptide sequences were plotted by position within the IgV_H_ sequence. The mass spectrometry proteomics data were deposited to the ProteomeXchange Consortium via the PRIDE^94^ partner repository with the dataset identifier PXD043144 and 10.6019/PXD043144.

### HLA expression plasmids

Expression plasmids for HLA-A*01:01 α-chain (pTCF5, Plasmid #180448) and β2m (pTCF158, Plasmid #180457) were purchased from Addgene. Single point mutations (V158A, R163T, or D166E) were introduced to this plasmid by site-directed PCR mutagenesis using complementary primers (Integrated DNA Technologies, Inc.) and Q5 High-Fidelity PCR Kit (New England Biolabs, E0555S). Plasmid DNA was sequence-verified for each HLA α-chain mutant.

### Recombinant HLA production

wildtype (WT) HLA-A*01:01 or single residue mutants of HLA-A*01:01 (V158A, R163T, D166E) were produced in-house as described.^95–97^ BL21 (DE3) *E. Coli* chemically competent cells were transfected with expression plasmids for human β2m, WT A*01:01 α-chain, or mutant A*01:01 α-chain. Expressed recombinant proteins were extracted from inclusion bodies and refolded by dilution of the α-chain and β2m into refolding buffer containing excess peptide (VTEHDTLLY, Biosynth Gardner, MA).^98^ The HLA WT and mutant complexes were purified by ion exchange chromatography (DEAE-C) and size-exclusion chromatography, buffer exchanged into PBS, concentrated, and stored at 4°C. For experiments requiring biotinylated HLA, the proteins were enzymatically biotinylated with BirA enzyme (Avidity), then purified by ion-exchange and size-exclusion chromatography.

### Generation of HLA-coated microbeads

Custom HLA cytometric bead arrays were generated as previously described^86^ using 4 µm and 5 µm carboxy functionalized array kits (Spherotech, PAK-4067-8K and PAK-5067-10K), conjugated with standard polymeric Streptavidin (SA, Southern Biotech), or monomeric streptavidin (monoSA, Sigma). SA or monoSA was buffer-exchanged using PD-10 columns (Cytiva) into PBS and diluted to 2 mg/mL. To conjugate SA or monoSA to unlabeled 4 and 5 µm Spherotech beads, the beads (1×10^8^) were isolated by centrifugation (3000g, 5min) and resuspended in 0.5 mL of 2 mg/mL of SA or mono SA. 0.5 mL of 6 mM 1-Ethyl-3-(3-dimethylaminopropyl)carbodiimide (EDC) in 0.05 M MES buffer pH 5.0 (Pierce) was added and the reaction mixture was rotated at RT overnight. Following incubation, the reaction was quenched with 0.1 mL of 1 M tris pH 8.0. The beads were washed twice in 1 mL PBS and resuspended at 1×10^8^/mL in PBS with 1% BSA and 0.005% NaN_3_ and stored at 4°C. Recombinant, biotinylated HLA (see above) was incubated with microbeads conjugated with either SA or monoSA (1×10^8^ microbeads/mL) at 1 mg/mL overnight at 4°C. The HLA-loaded beads were washed twice in 1 mL PBS and resuspended at 1×10^8^/mL in PBS with 1% BSA and 0.005% NaN_3_ and stored at 4°C until used. Standard SA-HLA coated beads were used in all assays except Ab inhibition studies (see below), which used monoSA-HLA coated beads to minimize non-specific inhibition due to HLA packing density.

### HLA cytometric bead arrays

Commercial (OneLambda, ThermoFisher) or custom (see above) HLA-coated microbeads were incubated for 15 min with analytes (single IgG1 rmAbs, plasma/serum, or appAbs) diluted in PBS or PBS with 1% BSA. Following incubation and washing, the beads were incubated with anti-IgG::PE or anti-IgG::AF488 (2.5 µg/mL, Southern Biotech) in PBS with 1% BSA for 15 min. Beads were washed, resuspended in PBS and analyzed on either a CytoFLEX Flow Cytometer (Beckman Coulter Life Sciences) or LSRFortessa (BD Biosciences). Data were analyzed in FlowJo v10.8.1 (BD). The gMFI of Ab bound to the different HLA allele-specific microbeads were calculated. Anti-influenza human IgG1 rmAbs^86^ were used as negative controls. The mean gMFI of the negative controls was subtracted from each allele-specific gMFI to obtain gMFI minus background (net gMFI). To select the minimum threshold for specific binding to HLA beads, the net gMFI of rmAb binding to three representative HLA alleles (A*01:01, A*02:01, and A*24:02) was compared to the L-SPR-calculated K_D_ of binding by the rmAb to each HLA allele. As 100% of tested rmAbs with an allele-specific net gMFI ≥40,000 had measurable allele-specific binding (K_D_ < 4.3 µM) by L-SPR, specific binding to the HLA beads was defined as a net gMFI ≥40,000.

### Binding of rmAbs to HLA A*01:01 mutants

MonoSA-conjugated microbeads that were loaded with WT A*01:01 or A*01:01 mutants were generated (see above), stained with analytes (rmAbs, appAbs, or sera/plasma) at multiple concentrations, and the gMFI of analyte binding to the beads displaying WT or mutant A*01:01 was determined. The half maximal effective concentration (EC_50_) of rmAb binding to each A*01:01 mutant was calculated using the sigmoidal, 4PL, x=concentration model in GraphPad Prism, with constraints (bottom = 0, top = 100). The gMFI of rmAb to A*01:01 mutant-coated beads was normalized to the maximum gMFI according to the following formula: % maximum gMFI = (gMFI of sample / highest gMFI for given protein) × 100%. To ensure accurate curve fitting, prozone points were omitted from the analysis. EC_50_ values were calculated from at least two technical replicates for each experiment. Serum anti-HLA Ab activity was interpolated from a standard curve using the pan-HLA Class I-reactive Ab W6/32 (Biolegend, 15.6 ng/mL to 2000 ng/mL) binding to WT A*01:01 and mutant HLA A*01:01 beads and corrected for dilution factor. The resulting standard curves were each fitted to a sigmoidal, 4PL, x=concentration curve. Each serum sample was initially assayed at a 1:500 dilution. In cases where the interpolated value for a given sample was outside of the vertical range of the standard curve, the serum sample was retested at a higher (1:4,000 or 1:12,000) dilution.

### Chimeric rmAb inhibition assays

Mouse IgG1 Fc chimeric (c) Abs (cE07 and cL02) were generated by fusing the human *IgV_H_* domain of E07 or L02 to the mouse IgG1 constant region. The human *IgV_L_* domain of E07 or L02 was then fused to either the mouse constant region kappa (for L02) or lambda (for E07) (Sino Biological) and the chimeric IgH and IgL proteins were co-expressed in 293 Freestyle cells. For Ab competitive inhibition analyses, monoSA-conjugated beads loaded with A*01:01 were produced (see above) then diluted in PBS alone (non-blocked beads) or diluted in PBS containing 5 μg/mL recombinant mouse chimeric cE07 or cL02 Abs (blocked beads) and then incubated for 10 min at RT. Analytes, including test human rmAbs, polyclonal appAbs, or polyclonal serum/plasma, were added at varying concentrations to the blocked and unblocked beads and bead suspensions were incubated for an additional 10 min. The beads were washed in PBS, stained for 10 min with anti-human IgG FC:PE (multi-species absorbed, Southern Biotech) diluted in PBS with 1% BSA, then washed and analyzed as described above. Percent inhibition of binding to the beads was calculated as follows: (unblocked gMFI – blocked gMFI)/(unblocked gMFI) × 100. All negative percent (%) values in analyses were set to 0. Percent inhibition values were calculated from at least two technical replicates for all experiments.

### X-ray crystallography and CryoEM sample preparation

HLA/Fab complexes were generated by mixing recombinant, non-biotinylated HLA-A*01:01 with either one Fab (1:1 molar ratio for E07/A*01:01, on ice) or with two Fabs (1:1:1 for L02/M07/A*01:01). Complexes were purified into PBS using size-exclusion chromatography on a Superdex S200 column. Complexes were concentrated and prepared for subsequent structural studies.

### X-ray crystallization, data collection, structure determination, and refinement

Purified E07/A*01:01 complexes were concentrated to 9.8 mg/mL and screened for crystallization conditions against commercially available kits using a Mosquito liquid handling robot (SPT Labtech). Crystallization conditions were optimized, and diffraction quality crystals were grown in 1.015 M sodium citrate tribasic dihydrate, 0.1 M sodium cacodylate pH 6.5 at a 1:1 ratio of protein to precipitating agent. Crystals were cryo-protected in crystallization solution supplemented with 20% (v/v) ethylene glycol and frozen. Diffraction data was collected at SER-CAT beamline 22-ID and processed with HKL2000.^99^ Data analysis was performed at UAB. Crystals exhibited a pseudo-cubic symmetry but had true C2 symmetry. Initial phases were solved by molecular replacement with PHASER^100^ using prior models of HLA (PDB ID: 6AT9),^101^ Ab domains: VH (PDB ID: 6PZH),^102^ VL (PDB ID: 4HK0),^103^ CH1 (PDB ID: 6AZZ).^104^ After identifying a trimeric assembly of HLA-Fab complexes, the assembly was used to find three additional copies of the assembly, yielding an asymmetric unit that contained 12 copies of HLA and 12 copies of the Fab. Models were iteratively rebuilt with COOT^105^ and refined with phenix.refine.^106^ Data collection and refinement statistics are provided (see Table S6). Though 12 copies of each protein could be identified in the asymmetric unit, domain α3 for some HLA and β2m had degenerate density following refinement due to high B-factors, likely a result of crystal packing, and were thus removed from the final model. After removal, electron density from Fo-Fc difference maps showed density for the additional domains, confirming their presence in the lattice. The final model contains six copies of the complete HLA-Fab complex, six additional copies of the Fab, three copies of the HLA domain with α3 removed, and two copies of β2m. Structure factors and final refined coordinates were deposited in the PDB (8T7R).

### CryoEM grid preparation and data collection

The L02/M07/A*01:01 complex was concentrated to 3.04 mg/mL in PBS. Grids for cryoEM were prepared by placing 3mL of solution on a 2/1-3T C-flat grid (R2/1 thick carbon, 300 mesh, Protochips), which was plunged into liquid ethane with a Mark IV Vitrobot. Grids were clipped and transferred to a G3 FEI-Krios electron microscope (La Jolla Institute for Immunology) equipped with a Gatan Bio-quantum energy filter with a 20eV slit. Images were detected on a K3 camera in counting mode (not super-resolution). The electron beam intensity was ∼15e^-^/pixel*s and 50 fractions were collected for every image, giving the specimen 1e/A*A*frame. The beam size was 900nm, allowing for collection of 6 images per hole in the sample.

### CryoEM data processing, model building and refinement

CryoEM data processing was performed by the UAB CryoEM Facility. The images were pre-processed with Warp^107^ to correct for movement, CTF parameters were estimated, and an initial set of particles was picked. CryoSPARC v4 was used for data processing.^108^ Pre-processing was repeated using patch motion correction and CTF estimation. From 14,572 exposures, blob picking and 2D classification yielded 93,263 preliminary particles for Topaz training (Figure S7A).^109^ Topaz autopicked 804,662 particles, which were initially downsampled to 2.64Å/pix and cleaned via 2D classification. The remaining particles were classified into four classes using ab-initio 3D reconstruction. This was done in quadruplicate for three iterations, keeping any particles that contributed to a good class. 252,746 particles from 3D classes corresponding to the L02/M07/A*01:01 complex were kept. After reverting to the original 0.66 Å/pix particles, these were refined using non-uniform refinement to a resolution of 3.14Å (Figure S7B-C).^110^ To initiate atomic model building, AA sequences were input to ColabFold via ChimeraX to generate AlphaFold predicted models.^111–113^ M07 and L02 were differentiated based on AAs with bulky side chains present in either, but not both, which could be clearly identified in the reconstruction. AlphaFold initial models were rigid-body fitted to the reconstruction using ChimeraX and refined in real space using Coot (Figure S7D).^105^ Phenix was used for global real-space refinement and model validation.^114^

### Structural analyses

Properties of Fab/HLA binding interfaces were analyzed using UCSF ChimeraX.^112^ Interacting residues at the Fab:HLA interface were identified using a heavy-atom (non-hydrogen atom) distance cutoff of ≤5 Å.^48^ Epitopic residues were defined as those with an interatomic distance between a heavy atom in the Fab and a heavy atom in the HLA complex ≤5 Å or any residues in the HLA complex with solvent-accessible surface area that was buried in the interface between the Fab and HLA-A*01:01. Hydrogen bonds were identified using UCSF ChimeraX hbond command with default criteria.^112^ *In silico* methods were used to observe interactions of Fab E07 with HLA complexes harboring mutations V158A, R163T, or D166E. Individual mutants were made with Coot. Complexes were subjected to the molecular dynamics energy minimization routine in YASARA.^115^

### Analysis of HLA residue properties

For consistency across structural models, a previously reported structure of HLA-A*01:01 (PDB: 3BO8)^116^ was used for calculations of HLA-A*01:01 residue solvent accessibility in the unbound state by PISA^117^ and Ellipro scores for residue protrusion.^118^ Physicochemical mismatches were compared as previously described^119^ for hydrophilicity and electrostatic mismatch, and AA substitution similarity scores were used for dissimilarity calculation.^120^ The AH-PBG region was defined as the HLA-A*01:01 α-chain α1 domain residues 50-85 and α2 domain residues 138-175, which frame the peptide-binding groove (PBG).^53^ To calculate residue distances from the center of the PBG, the previously reported HLA-A*01:01 (PDB: 3BO8)^116^ structure was used and the center of the PBG was defined as the α-carbon of the 5^th^ residue (T5) in the peptide nonamer (EADPTGHSY) presented within the PBG. Distances were calculated between the peptide T5 residue α-carbon and the α-carbon of each AA in A*01:01 using UCSF ChimeraX.^112^

### B cell epitope prediction using computational tools

B cell epitopes within the HLA-A*01:01 protein were predicted using GraphBepi^58^ and DiscoTope-3.0.^59^ For both tools, the previously reported HLA-A*01:01 (PDB: 3BO8)^116^ structure was used. For GraphBepi, default settings were used. For DiscoTope-3.0, the epitope confidence threshold was set to 1.50.

### HLA genotyped cohort transplant simulation

We examined published^121^ HLA genotyping data derived from a cohort of 310 United States subjects and selected 247 subjects who did not express an A*01:01 self-allele. The AA residues that were mismatched between the self-HLA-A alleles expressed by each individual and the non-self A*01:01 protein were identified and the total number of solvent-accessible residues that were classified as 2MM and located within the AH-PBG region was determined for each subject. The percentage of these 2MMs that were contained within the E07 and L02 epitopes was calculated. Residues were classified as solvent-accessible if the residue non-buried surface area was 250 Å^2^.

### Statistical analyses

All statistical comparisons were performed in Prism (GraphPad Software). Comparisons of an observed distribution with the expected distribution were performed using the binomial test and the confidence interval of the proportion was calculated using the hybrid Wilson/Brown method in Prism. Comparisons of proportions were made using Fisher’s exact test. Following confirmation of distribution normality by the Shapiro-Wilk test, comparisons of group means was made using an unpaired t-test. If distributions were assessed to be non-normal, group ranks were compared via the Mann-Whitney test. Paired analyses were performed using the Wilcoxon matched-pairs signed rank test. Comparisons of multiple groups were performed using the Kruskal-Wallis test with Dunn’s multiple comparisons test (nonparametric with no matching), the Friedman test with Dunn’s multiple comparisons (nonparametric with matching), or the Brown-Forsythe and Welch ANOVA tests with Dunnett’s T3 multiple comparisons test (parametric with no matching). Spearman correlations were used to assess association between two nonparametric variables. Hierarchical clustering, heatmap plots, and other visualizations were created using R v4.2.0 (R Core Team 2022),^122^ Matlab (R2020a, The Mathworks Inc., Natick, MA), UCSF ChimeraX v1.5,^112^ Cytoscape v3.8.2,^91^ FlowJo v10.8.1 (BD, Franklin Lakes, NJ), Prism v10.2.1 (GraphPad Software, LLC., Boston, MA), or Biorender.com (Toronto, Canada).

### Data and Code Availability

IgV_H_ sequencing data has been deposited on GEO (GSE235533). CryoEM data has been deposited in PDB, EMDB, and EMPIAR (8T6M, EMD-41072, EMPIAR-11591). X-ray crystallography data has been deposited in PDB (8T7R). Proteomics data is deposited in ProteomXchange (PXD043144). Original code for the plasma proteomics analysis is deposited on Zenodo (10.5281/zenodo.8039145).

